# Examination of digital images from Macaulay Library to determine avian molt strategies: A case study on molts and plumages in eight species of North American hummingbirds

**DOI:** 10.1101/2021.02.03.429637

**Authors:** Peter Pyle

## Abstract

I examined a total of 27,581 images of 6.345 individuals from the Cornell Lab of Ornithology’s Macaulay Library to clarify conflicting reports on molt and plumage strategies in eight species of hummingbirds that breed or have bred primarily in the southwestern United States. Fixed replacement sequences from two nodes among primaries and two nodes among secondaries were found without exception, conforming to the findings of previous studies. I concluded that the preformative molt is limited to partial in three species, partial to incomplete in three species, partial to complete in one species, and complete in one species. These molt strategies could be interpreted as having differentiated through synapomorphy, with species between currently recognized clades varying in the extent of their preformative molts; however, given the plastic nature of molt strategies, I predict that this variation will be shaped more by environmental factors than by synapomorphy. Results of this study additionally clarify molt terminology in Trochilidae as based on homologies and establish new criteria for age determinations. The Macaulay Library clearly provides an important resource for the investigation of avian molts and plumages. The results of a validation exercise that I conducted indicate that banders and field ornithologists with a wide range of previous experience can collect accurate data in this manner. I present a road map for such studies and suggest many other questions on avian molt that can also be investigated, including how timing of molts vary geographically and by habitat and how remigial replacement sequences proceed in little-known bird families. I encourage contributors to the Macaulay Library to take and upload images of birds in molt or in worn plumages.

## Introduction

Our understanding of avian molt strategies has lagged behind that of other aspects of avian natural history (Bridge 2011, Marra et al. 2015), and this lack of knowledge is especially acute among the large number of bird species found in equatorial regions (Craig 1983, Mulyani et al. 2017, Johnson and Wolfe 2018). Although study of specimens has been instrumental in advancing our knowledge of avian molts, relatively few birds have been collected while undergoing active molt (Rohwer et al. 2005), and large sample sizes are often needed to fully document variation in timing, location, and extent of molts within a species’ annual cycle and throughout its geographic range.

Traditionally, hummingbirds in the United States and elsewhere were assumed to undergo complete preformative and prebasic molts and to lack prealternate molts (Williamson 1956, Baltosser 1995, Pyle 1997, Howell 2002, Wolfe et al. 2009). However, the discovery of definitive prealternate molts in Ruby-throated *(Archilochus colubris)* and Rufous *(Selasphorus rufus)* hummingbirds has lead to other proposed terminologies (Dittmann and Cardiff 2009, Howell 2010, Weidensaul et al. 2020), including a strategy that considers preformative molts in these species to be partial (Sieburth and Pyle 2018). With the exception of the presence or absence of prealternate molts, the strategies of the eight species in genera *Archilochus, Calypte,* and *Selasphorus* that breed in the United States (hereafter “northern” hummingbirds) are reasonably well documented (Williamson 1956, Baltosser 1995, Pyle 1997, Pyle et al. 1997, Howell 2002, Williamson 2002). However, those of the eight species of genera *Eugenes, Lamphornis, Calothrax, Cynanthus, Basilinna, Leucolia, Saucerottia,* and *Amazilia,* that breed or have bred primarily in Texas and the southwestern United States (hereafter “southwestern” species), are not as well known. Most of these species have ranges that extend to southern Mexico or Central America, where geographic variation in seasonal regimes and life-history requirements may complicate molt strategies.

Previous authors (e.g., Pyle 1997) attempted to confirm reports in the literature on hummingbird molt by examining specimens and data from banding stations. For the eight northern species generally there have been adequate sample sizes from these sources to accurately assess molt strategies, including of specimens collected on winter grounds in Mexico (Pyle et al. 1997, Sieburth and Pyle 2018). However, for the eight southwestern species, sample sizes of specimens and captured birds have often been sufficiently lacking to gain a full understanding of strategies. Currently there is conflicting information on timing and extents of molts in these species as presented by Pyle (1997), Howell (2002), Williamson (2002), the Birds of the World accounts (Billerman et al. 2020), and additional data collected from banding stations in the United States and Mexico (Wethington 2020).

Beginning in the mid-2000s, the advancement of digital technology has allowed detailed examination of feathers and feather tracts in images of birds, which in turn has been used to study molts and plumages (Pyle 2008a, Viera et al. 2017, Panter 2021). Since this time, the quantity of available on-line images has increased exponentially, expanding the potential to augment data on bird molt collected from specimens. The Cornell Lab of Ornithology’s Macaulay Library archives audio and video recordings and images of birds and other wildlife for scientific research, education, and conservation. Virtually all of the bird images archived at the library were contributed as part of eBird, a citizen science project allowing both birders and researchers to archive count data, images, and other media resulting from observations in the field (Sullivan et al. 2009). eBird provides comprehensive search functions of the Macaulay Library that allows viewing of digital images after applying various filters including location(s), year(s) and month(s) of observation. Images can be ordered by date of observation, date uploaded, or a quality rating from users. Currently there are over 20 million images of 10,056 bird species had been contributed to the library (M. Medler pers. comm.), typically representing images from throughout a species’ annual cycle, and providing a tremendous resource for the study of avian plumages and molts.

I examined images archived at Macaulay Library to better document and clear up inconsistent information on molts and plumages for the eight southwestern hummingbird species. My goals included assessing the extent of the preformative molt (partial, incomplete, or complete), establishing timing for all molts and plumages, evaluating replacement sequences among flight feathers, and applying results to the accurate ageing and sexing of these eight species and to our understanding of the evolution of molts, hence, molt terminology in these and other hummingbirds (Humphrey and Parkes 1959, Howell et al. 2003, Sieburth and Pyle 2018). I also undertook a validation study with banders and field ornithologists to test the applicability of this methodology. My primary goal is to provide a case study for using the Macaulay Library to study avian molts around the world.

## Methods

Species examined for this analysis were Rivoli’s Hummingbird *(Eugenes fulgens),* Blue-throated Mountain-gem *(Lamphornis clemenciae),* and Lucifer *(Calothrax lucifer),* Broad-billed *(Cynanthus latirostris),* White-eared *(Basilinna leucotis),* Violet-crowned *(Leucolia violiceps),* Berylline (*Saucerottia beryllina*), and Buff-bellied (*Amazilia yucatanensis*) hummingbirds. I sought to assess molt patterns within populations of these species that breed or occur north of Mexico. Therefore, I set eBird’s location filter of Macaulay Library images to the United States. For each species I used the month filter to examine images for each month of the year. For Lucifer, White-eared, and Berylline hummingbirds, I concluded that sample sizes of images from the United States year-round were insufficient to gain an accurate assessment of molt patterns. I therefore set the filter to Mexico and augmented the sample by examining images taken in the northern tier of Mexican states and those on the Mexican Plateau south to the Distrito Federal, with the assumption that these bioregions included wintering individuals from the United States or breeding populations that exhibited similar molt strategies. Within each month I ordered the images by date, from oldest to newest. This allowed better tracking of individual hummingbirds, for example, those at popular feeding stations, thereby minimizing duplication of data from the same individuals.

All images at Macaulay Library of these eight species taken in the United States and uploaded through July 2020 were reviewed. Hummingbirds misidentified to species (< 1%) were excluded. Data were recorded only from images that could be properly assessed for both plumage (age) and molt status; e.g., all primaries of the wing were visible or accounted for in molt (Figure 1). In many cases the eBird Checklist contained multiple images of the same individual, which helped with accurate determinations. Individuals that were not confidently aged were excluded. I also excluded images of the same individual within a month as conservatively as possible based on molt and plumage status, date, location, eBird checklist data, age, appearance, and bill pattern. Generally, a bird of similar molt status, plumage, and appearance within 7 d of a previous observation at the same location was assumed to be the same individual. Individuals with images that spanned months were recorded for each month of occurrence.

**Figure 1.**
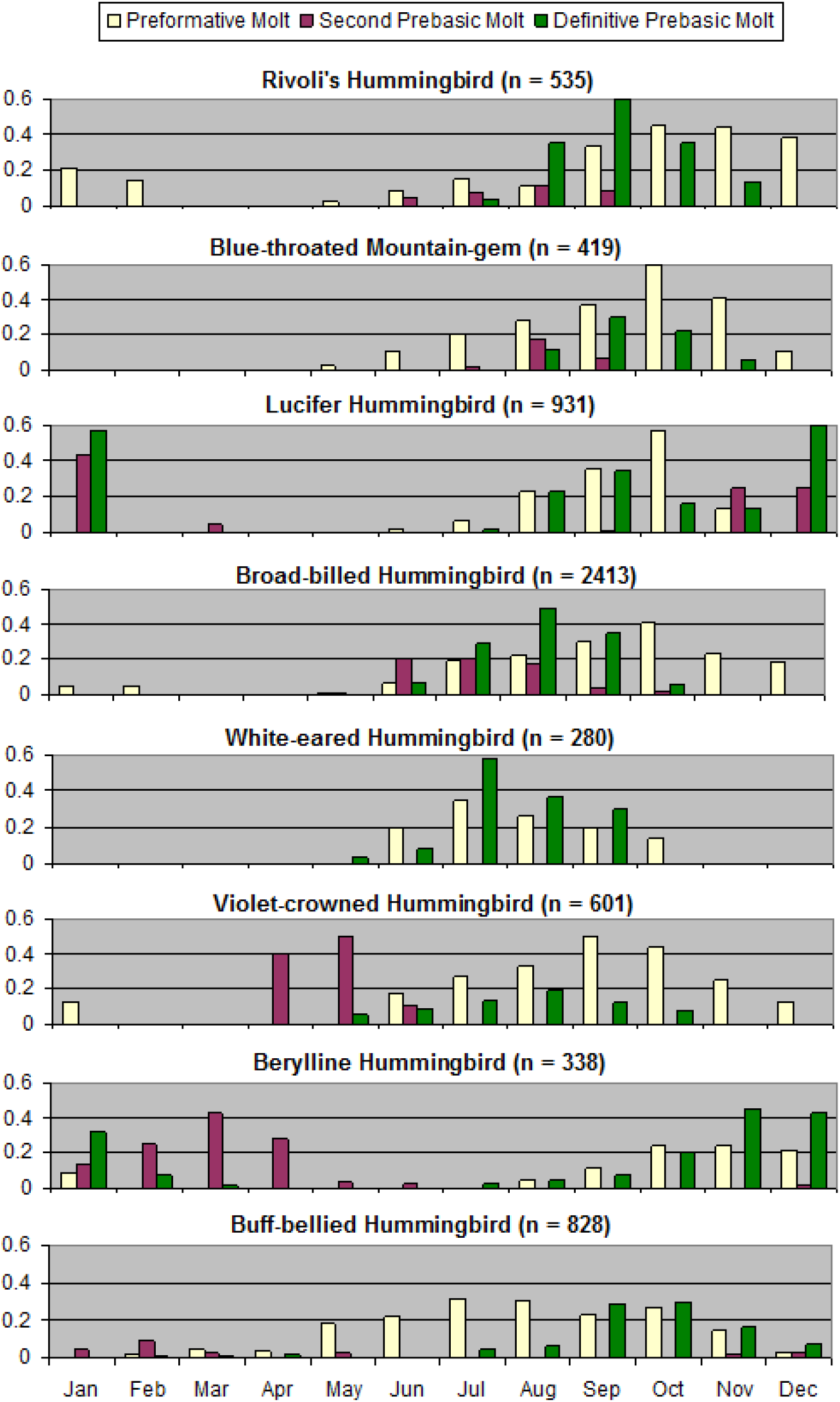
The timing of molt in eight species of hummingbirds that breed in the southwestern United States. Bars represent proportion of the entire monthly sample that were undergoing each molt; see Supplemental Material Table S1 for specific sample sizes for each species by month.

For each individual I determined plumage and molt status. Plumages in both males and females were identified following the ageing criteria of Pyle (1997), Howell (2002), and Williamson (2002), along with new criteria presented here (Supplemental Figures S1-S9). Criteria based on wing feathers, rectrices, and for some species bill color were emphasized; that of iridescent feathering in males was evaluated with caution due to effects of lighting on the perceived coloration of these feathers in digital images. Extent of corrugation at the base of the culmen (Ortiz-Crespo 1972, Yanega et al. 1997, Pyle 1997) was also examined but could only be evaluated on a small proportion of images. For individuals in active molt, replacement sequence of primaries, secondaries, and rectrices was assessed (Figure 1). Primaries were numbered proximally from p1 (inner) to p10 (outer) and secondaries distally, from s1 (outer) to s6 (inner). Comparison of primary and secondary spacing (morphology) in images of birds not in active molt was employed to help determine precise sequences in molting birds, and symmetry among new, molting, and old feathers within both wings was confirmed, when possible, to ensure that missing feathers reflected molt. Arrested or suspended molts among flight feathers (cf. Pyle et al. 1997) were noted as contrastingly new feathers in sequence among older unreplaced feathers. Molt and molt limits among body feathers and upperwing secondary coverts were also assessed by evaluating pin and growing feathers along with contrasts between new and old feather generations. See the Supplemental Materials File for more detail on this methodology.

I categorized each individual into one of six plumage or molt states: 1) juvenile plumage (prior to evidence of preformative molt), 2) undergoing preformative molt, 3) formative plumage of body or flight feathers, 4) undergoing second prebasic molt of flight feathers, 5) definitive basic plumage, and 6) undergoing definitive prebasic molt of flight feathers. Partial preformative molts (excluding primaries) are often protracted and/or suspended resulting in less-precise assignment of preformative molt or formative plumage. To categorize these I looked for pin and growing feathers and also assessed when development of definitive-like appearance appeared to culminate within the entire sample of first-cycle males, including long-staying individuals undergoing and completing preformative molt. Timing of molts and plumages in hummingbirds, except for gorget-feather replacement in males, shows little variation by sex (Williamson 1956, Pyle et al. 1997, Sieburth and Pyle 2018), and this also accorded with exploratory examination of Macaulay Library data for this study. Therefore, counts included both sexes combined. Images of interest are referenced by their Macaulay Library identifiers (“ML” followed by 8 or 9 numerals) and in some cases eBird Checklist identifiers (“S” followed by 7 or 8 numerals) when multiple images of the same bird documented the point of reference.

Examination of images to study avian molt may require extensive previous experience with captured birds or specimens. To test whether or not banders and field ornithologists with a varying range of previous field experience can collect accurate data on molt from images, I circulated a validation study which included images of 11 hummingbirds from the Macaulay Library (see Supplemental Materials File). Each participant was asked to evaluate their previous experience with banding and field ornithology, to determine the age of the individual, to score the status of molt (active or inactive), to score the condition of each of the 10 primaries (new, growing, missing, or old), and to record the number of minutes it took to age and score each individual. Participants were given Supplemental Figures 1-8 as a study guide before undertaking the exercise.

## Results

A total of 27,581 images from the Macaulay Library of the eight southwestern hummingbirds was examined for this study (Supplemental Table S1). These included 6,345 individuals from images that were of sufficient quality to assess plumage (age) and molt status. Total individuals by species ranged from 280 White-eared Hummingbirds to 2,413 Broad-billed Hummingbirds, totals by month ranged from 248 individuals for February to 1,245 for August, and totals by species in a month ranged from 4 Lucifer Hummingbirds in February to 639 Broad-billed Hummingbirds in July (Supplemental Table S1). Samples of > 25 individuals were recorded for 75% of the months by species.

Sequence of feather replacement among primaries consistently proceeded from a node at p1 distally and a node at p10 proximally, with p9 being the last primary replaced (Figure 1). Among images of 1,373 individuals recorded undergoing active primary molt, no exceptions to this sequence were observed (cf. Supplemental Figures S1-S8), including among >10 known individuals that could be tracked for all or large portions of the molting period. The six secondaries of these species began to be replaced when p6 had dropped (e.g., ML 181183661, [ML46645931](https://macaulaylibrary.org/asset/46645931), [ML122879681](https://macaulaylibrary.org/asset/122879681)). Among 71 individuals in which active secondary molt could be evaluated, replacement invariably proceeded proximally from a node at the innermost feather s6 and distally from a node at the outermost feather s1 (Figure 1). The orders in which s1 and s6 and s3 and s5 were molted were variable, but s4 was always the last feather to be replaced, near to or following completion of primary molt (e.g., ML 195887161, [ML34535671](https://macaulaylibrary.org/asset/34535671), [ML100323371](https://macaulaylibrary.org/asset/100323371), ML 33989911). Sequence of rectrix replacement was more difficult to evaluate in images but typically began with the central rectrices when p7 or p8 were dropped (e.g., [ML184013981](https://macaulaylibrary.org/asset/184013981), ML 188765281, [ML34654481](https://macaulaylibrary.org/asset/34654481)), after which replacement of remaining rectrices generally proceeded rapidly and distally (e.g., [ML238319331](https://macaulaylibrary.org/asset/238319331), S56405130), with the outermost (r5) often replaced before r4 and/or r3 (e.g., [ML238400031](https://macaulaylibrary.org/asset/238400031)).

Suspended or arrested molts among non-molting remiges were rare, being recorded in only 23 individuals (< 0.01% of 3,652 non-molting hummingbirds), of six species, Rivoli’s Hummingbird, Blue-throated Mountain-gem, and Broad-billed, White-eared, Violet-crowned, and Buff-bellied hummingbirds. These were recorded during both preformative molts (see below) and definitive prebasic molts (e.g., [ML45716111](https://macaulaylibrary.org/asset/45716111), [ML212853441](https://macaulaylibrary.org/asset/212853441), ML 86250771, [ML51351041](https://macaulaylibrary.org/asset/51351041), S41676864), including individuals that had replaced all remiges except for the s4 (e.g., [ML48860261](https://macaulaylibrary.org/asset/48860261), [ML42333141](https://macaulaylibrary.org/asset/42333141), S2611244). Suspended or arrested molts were recorded at a single location within the above sequences, with the exception of one Buff-bellied Hummingbird that had suspended molt after replacing p1 and p6 most recently (ML 22821451), perhaps following an earlier arrested molt. No retained rectrices resulting from suspended or arrested molts were noted but these could easily have been missed.

Sample sizes for the six molt and plumage categories, by month, for the eight species are given in Supplemental Table S1. Dates for juveniles ranged from 2 February in Buff-bellied Hummingbird to 18 November in Berylline Hummingbird, and temporal duration periods for recorded juveniles ranged from 78 d in Violet-crowned Hummingbird to 184 d in Rivoli’s Hummingbird (Supplemental Table S2). The preformative molt was first detected from nine (in Berylline Hummingbird) to 80 (in White-eared Hummingbird) days following the earliest juveniles recorded in the spring. Temporal periods for the preformative molt among populations ranged from about 5 mo in White-eared Hummingbird, Lucifer Hummingbird, and Berylline Hummingbird, to about 7 mo in Blue-throated Mountain-gem and Violet-crowned Hummingbird, about 8 mo in Broad-billed Hummingbird, 9.4 mo in Rivoli’s Hummingbird (282 d), to 10.3 mo in Buff-bellied Hummingbird (Figure 2, Supplemental Table S2).

**Figure 2.**
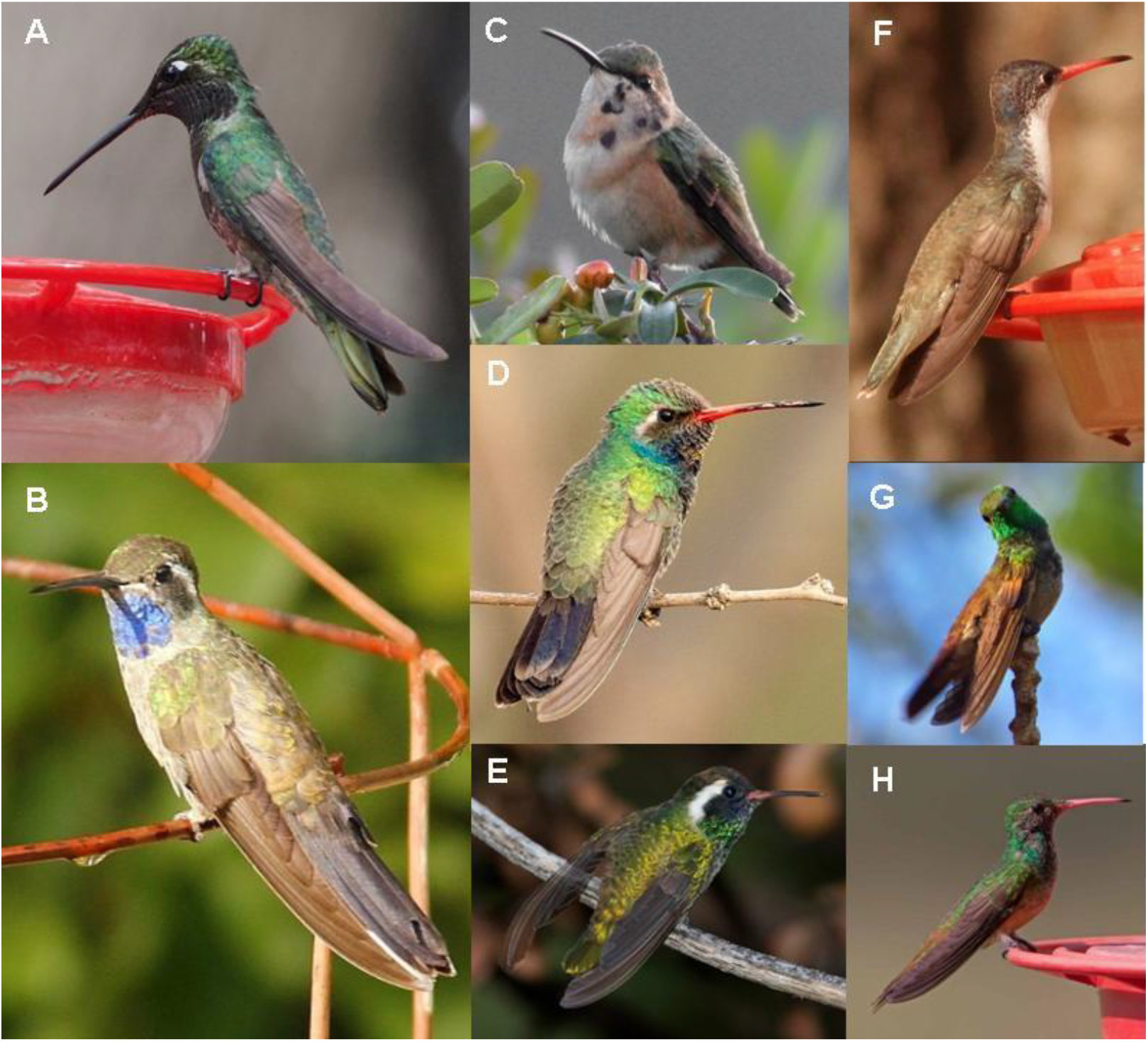
Examples of formative plumage in eight species of hummingbirds that breed in the southwestern United States. (**A**) Rivoli’s Hummingbird *(Eugenes fulgens),* 4 Aug 2019; (**B**) Blue-throated Mountain-gem *(Lamphornis clemenciae),* 4 Aug 2012; (**C**) Lucifer Hummingbird *(Calothrax lucifer),* 5 Oct 2009; (**D**) Broad-billed Hummingbird *(Cynanthus latirostris),* 2 May 2019; (**E**) White-eared Hummingbird *(Basilinna leucotis),* 5 Aug 2008: (**F**) Violet-crowned Hummingbird *(Leucolia violiceps),* 6 Mar 2016; (**G**) Berylline Hummingbird *(Saucerottia beryllina),* 11 Feb 2017; and (**H**) Buff-bellied Hummingbird *(Amazilia yucatanensis*), 26 Apr 2017. Except for White-eared Hummingbird, note the retained juvenile primaries, worn brown secondaries, and molt limits among upperwing secondary coverts in most or all images. The White-eared Hummingbird (**E**) is finalizing a complete preformative molt (aged by dull bill color) after which formative plumage resembles definitive basic plumage in appearance. Photos cropped for enlarged presentation and used by license agreement from the Macaulay Library © Gjon Hazard (**A**, [ML171639201](https://macaulaylibrary.org/asset/171639201)), Ken Murphy (**B**, [ML53554351](https://macaulaylibrary.org/asset/53554351)), Ed Thomas (**C**, [ML168356961](https://macaulaylibrary.org/asset/168356961#_ga=2.221619032.1360707014.1598479728-1260102835.1597029967)), Philip Kline (**D**, [ML156749041](https://macaulaylibrary.org/asset/156749041)), Bill Hubick (**E**, [ML188765291](https://macaulaylibrary.org/asset/188765291)), Debby Parker (**F**, [ML25520031](https://macaulaylibrary.org/asset/25520031)), William Proebsting (**G**, [ML49162901](https://macaulaylibrary.org/asset/49162901)), and Joshua Covill (**H**, [ML56665951](https://macaulaylibrary.org/asset/56665951)).

I concluded that the preformative molt is typically limited to partial in three species, Lucifer, Berylline, and Buff-bellied hummingbirds (Figure 3 and Supplemental Figures S3, S7, and S8). These species replace variable amounts of body feathers and upperwing secondary coverts, from a few body feathers only to most or all body feathers and secondary coverts, but replaced no primaries, primary coverts, secondaries, or rectrices until commencement of the second prebasic molt. Most Rivoli’s Hummingbirds, Blue-throated Mountain-gems, and Violet-crowned Hummingbirds also undergo partial preformative molts (Figures 1 and Supplemental Figures S1, S2, and S6), although small proportions, one of 126 first-cycle Rivoli’s Hummingbirds in October-August (0.8%), one of 95 first-cycle Blue-throated Mountain-gems in November-August (1.1%), and three of 140 first-cycle Violet-crowned Hummingbird in August-May (2.1%) were replacing or had replaced two to six inner primaries during what I judged to be incomplete preformative molts (Figure 4).

**Figure 3.**
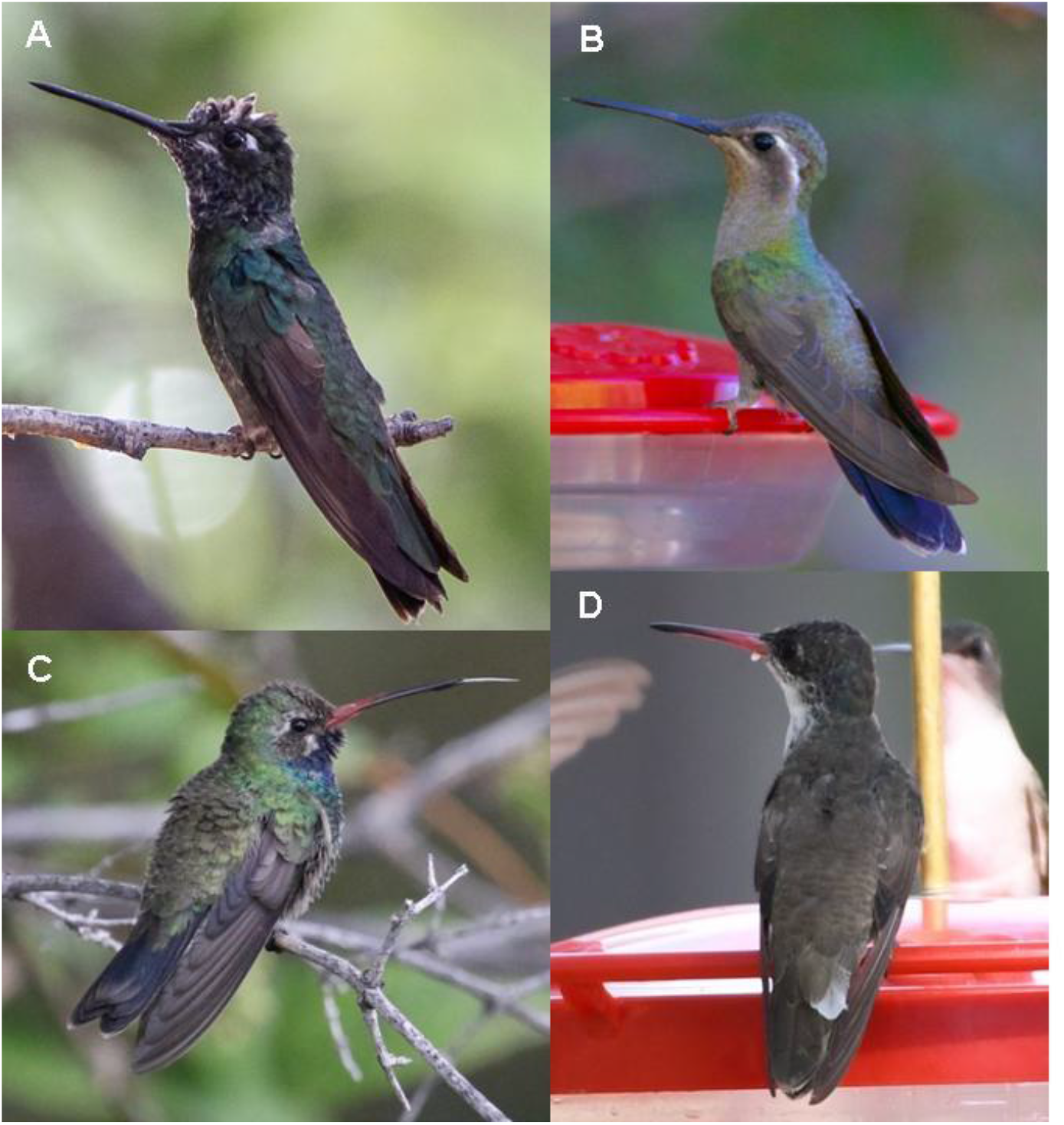
Incomplete (arrested or suspended) preformative molts in four species of hummingbirds that breed in the southwestern United States. (**A**) Rivoli’s Hummingbird *(Eugenes fulgens),* 26 May 2018, having replaced p1-p3; (**B**) Blue-throated Mountain-gem *(Lamphornis clemenciae),* 4 Aug 2012 having replaced p1-p4; (**C**) Broad-billed Hummingbird *(Cynanthus latirostris),* 16 September 2019, having replaced p1-p4; and (**D**) Violet-crowned Hummingbird *(Leucolia violiceps),* 31 Aug 2019 replacing p1-p6. Photos cropped for enlarged presentation and used by license agreement from the Macaulay Library © Lydie Mason Warner (**A**, S46054488), Gordon Atkins (**B**, [ML101073691](https://macaulaylibrary.org/asset/101073691)), Russ Morgan (**C**, S59856736), and Max Leibowitz (**D**, S59412857).

For Broad-billed Hummingbird, the preformative molt varied from partial to complete (Figures 2 and 3 and Supplemental Figures S4). Active molting of primaries and rectrices during the preformative molt of this species (*n* = 29) was recorded from 3 August ([ML86966331](https://macaulaylibrary.org/asset/86966331)) to 25 January (S33932855) with active molting of secondaries recorded through 16 February ([ML208941611](https://macaulaylibrary.org/asset/208941611)). In November-December, 56% of 32 first-cycle birds were molting or had molted primaries, and in December-May at least 10 of 222 first-cycle individuals (4.5%) had suspended or arrested primary molt, most often at p2 (e.g., [ML77579011](https://macaulaylibrary.org/asset/77579011)) or p4 (Figure 4; see also, e.g., [ML47852161](https://macaulaylibrary.org/asset/47852161), [ML22932271](https://macaulaylibrary.org/asset/22932271)). Some males underwent a complete preformative molt of flight feathers but did not acquire definitive appearance of body plumage whereas others acquired complete or near-complete definitive appearance in body feathering but retained juvenile flight feathers (Figure 2 and Supplemental Figures S4 and S9). Some Broad-billed Hummingbirds following complete molts likely become indistinguishable from individuals in definitive basic plumage, and were categorized as in definitive basic plumage here.

For White-eared Hummingbird I concluded that the preformative molt was complete. It was the only one of the eight species in which timing of preformative and later molts was similar, the replacement of primaries commencing at the same time or before juvenile body feathers began molting (Supplemental Figure S5; [ML252080431](https://macaulaylibrary.org/asset/252080431#_ga=2.204976787.2056923685.1595870569-1093485893.1549930479), S11291169) and completing following body-feather replacement, at which time males had acquired definitive-like appearance (Figures 1 and 2 and Supplemental Figure S5). It was also the only species in which no males following the preformative molt showed predefinitive appearance (*n* = 64). The longer period for juveniles recorded for this species (80 d) than the others (10-57 d) may also relate to the complete molt, juvenile feathers not needing to last for five months or more. As a result of this complete preformative molt, White-eared Hummingbirds in formative vs. definitive basic plumage and undergoing the second vs. definitive prebasic molts could not be distinguished in images for this study, with the exception of some in formative plumage with dull red bill colors.

Formative plumage in males (and in some cases females) of these seven species, as aged by flight-feather characteristics, generally did not reach definitive appearance of body feathering, varying from showing no or a few iridescent display feathers in male Lucifer Hummingbirds to showing nearly full to full definitive appearance in male Broad-billed, Berylline, and Buff-bellied hummingbirds (Figure 2 and Supplemental Figures S1-S4, S6-S9). Formative plumages in male Rivoli’s Hummingbird and Blue-throated Mountain-gem, and in both sexes of Violet-crowned and Buff-bellied hummingbirds, were variable and intermediate but few birds in formative plumage appeared to have acquired definitive appearance of body feathering (Supplemental Figures S1, S2, S6, and S9). By contrast, definitive basic males of all eight species (as aged by flight-feather characteristics) showed full definitive appearance, with the exception of a small proportion of Rivoli’s Hummingbirds that had small and variable amounts of brown feathering in the lower breast; further study is needed on whether or not this may represent second basic plumage.

Within the populations, the temporal duration period for the second prebasic molt ranged from 63 d in Blue-throated Mountain-gem and 65 d in Violet-crowned Hummingbird to 188 days in Buff-bellied Hummingbird, and for the definitive prebasic molt duration ranged from 94 d in Blue-throated Mountain-gem to 271 d in Buff-bellied Hummingbird (Supplemental Table S2). With the exception of Buff-bellied Hummingbird the seasonal timing for these molts was well defined (Figure 2 and Supplemental Table S2). Known individual hummingbirds take less time within these periods to molt; e.g., a Berylline Hummingbird in Arizona in 2020 was documented completing a definitive prebasic molt in 49 d, from dropping p1-p3 of 26 April (S67876574) to completing growth of p9 and s4 on 14 June ([ML244918451](https://macaulaylibrary.org/asset/244918451)), and a Buff-bellied Hummingbird in Florida that had dropped inner primaries on 4 November 2016 ([ML39279451](https://macaulaylibrary.org/asset/39279451)) was completing molt 69 d later on 28 January 2017 ([ML46926141](https://macaulaylibrary.org/asset/46926141)). Timing of the second prebasic molt differed but overlapped that of the definitive prebasic molt in all seven species, the overlap being earlier than the definitive prebasic molt in Rivoli’s Hummingbird, Blue-throated Mountain-gem, Broad-billed Hummingbird, and Violet-crowned Hummingbird, and later than the definitive prebasic molt in Lucifer, Berylline, and Buff-bellied hummingbirds (Figure 2). Based on my conclusions on the evolution of these molts, however, the second prebasic molt occurred earlier in timing than the definitive prebasic molt in all seven species (see Discussion).

Seventeen field biologists (including the author) participated in a validation exercise to ensure that this methodology can be used to collect accurate data on molt (Supplemental Materials File). Participants correctly aged the 11 hummingbirds (first-year or older) 83% of the time, reached a correct conclusion on molt status (active or inactive) 93% of the time, and provided correct answers for the condition of each primary (new, missing, growing, or old) from 83% to 91% of the time. For individual primary cells, correct answers ranged from 6% and 18%, up to 100% for most, with a mean of 87.2%. The mean time it took to age and score each hummingbird was 3.7 minutes. The mean proportion of correct answers for the 132 cells was 87.5%, ranging from 80.3% to 95.4% among the 17 observers. Among participants with low, medium, and high experience levels, correct answers were provided for 83.1%, 87.1%, and 88.9% regarding banding experience and 87.6%, 86.6%, and 88.9% regarding field experience, respectively.

## Discussion

Sequence of primary molt in hummingbirds has previously been reported to be distal from a node at p1, proximal from a node at p10, and with p9 the last feather replaced (Wagner 1955, Williamson 1956, Stiles 1995, Pyle 1997, Howell 2002). This sequence was confirmed with few or no exceptions among 1,373 molting hummingbirds of all eight species in this study. Results of this study also indicate replacement nodes among secondaries to be fixed, with proximal replacement from s1 and distal replacement from s6 resulting in s4 being the last secondary replaced, without exception within my sample, including for Lucifer Hummingbirds (e.g., [ML79210961](https://macaulaylibrary.org/asset/79210961)), contradicting reports by Wagner (1955) of replacement from nodes in the center of the tract (see also Stiles 1995). These remegial replacement nodes and directions are consistent with those found by Williamson (1956) for Anna’s Hummingbird *(Calypte anna)* and by Stiles (1995) for 13 hummingbird species in Costa Rica, although Stiles also found that the last secondary replaced was s3 or s5 rather than s4 in a small proportion (6.2%) of 242 individuals in his study. It is possible that variable sequences may follow arrested molts, which appear to be more common in species of tropical rather than in temperate habitats (Pyle et al. 2016), perhaps including Buff-bellied Hummingbird in this study.

Unlike timing, location, and extent of molts, sequential replacement of remiges in birds appears very fixed (cf. Pyle 2013), in which case I predict that these four remigial nodes and replacement directions will be found in all hummingbird species. Precise sequence among different replacement waves (e.g., in hummingbirds, initiation at either s1 or s6 or order of s3 vs. s5) and terminal feathers where waves converge is less fixed, evolutionarily, and may vary in birds according to wing physiology, flight requirements, or other parameters. My results on rectrix sequence also comport with those of Stiles (1995). The p9 is the longest primary in hummingbirds and it has been proposed that its replacement follows that of p10 to maintain wing integrity in a bird family that relies heavily on flight for existence (Greenewalt 1975, Stiles 1995). A similar sequence among primaries in family Ardeidae (Shugart and Rohwer 1996, Pyle 2008b) has evolved independently, perhaps for different reasons.

Additional results of this study otherwise clarify molt strategies in these eight southwestern hummingbirds to a substantial degree. For example, preformative molts in seven species are here interpreted to be partial in most individuals, differing from previous interpretations that they were complete (Pyle 1997, Howell 2002). In three of these species, Rivoli’s Hummingbird, Blue-throated Mountain-gem, and Broad-billed Hummingbird, juvenile primaries can be retained for close to a year, consistent with strategies in most other birds with partial preformative molts (Howell et al. 2003; Pyle 1997, 2008b, Jenni and Winkler 2020). Lucifer, Berylline, and Buff-bellied hummingbirds have molts more similar to northern North American species, in which body feathers are partially replaced during a preformative molt well before primaries are replaced as part of the second prebasic molt (see below). The timing of the second prebasic molt of Violet-crowned Hummingbird appears to be intermediate between these two groups and indicates that they may not breed in their first spring, although its apparently short duration may allow them to breed later in summer, following the molt. The extent of preformative molt in four species, Rivoli’s Hummingbird, Blue-throated Mountain-gem, Broad-billed Hummingbird, and Violet-crowned Hummingbird can at least occasionally include primaries and in White-eared Hummingbird it is complete. Variation in preformative molt extent, from partial to incomplete to complete, has also been documented within other bird species and genera, such as those among Scolopacidae, Tyrannidae, Fringillidae, Passerellidae, and Cardinalidae (Pyle 1997, 2008b), and perhaps is correlated with habitat use and extent of solar exposure on an annual basis (Pyle 1998, 2008b, Guallar et al. 2020). White-eared Hummingbird is the smallest of the eight species treated here (Billerman 2020), and this could also be a factor in its undergoing a complete preformative molt, as extent of partial or incomplete molts in birds generally increases with decreasing body size (Kiat and Izhaki 2016).

Results of this study also help clarify previous discrepancies on timing of complete molts in these eight southwestern hummingbird species. For example, in Broad-billed Hummingbird, Pyle (1997) reported that populations in the United States underwent the first molt of primaries in November-May and definitive prebasic molts in October-April; Howell (2002) concluded that the definitive prebasic molt commenced in April-September and completed in July-January, with first molt of primaries averaging later in timing; and Williamson (2002) indicated that the definitive prebasic molt occurred in May-September and the first molt of primaries occurred in July-November of the same year. Results of this study, by contrast, indicate that some birds first replace primaries during the prefomative molt in August-January, others replace them at the second prebasic molt in May-September of the following year, and the definitive prebasic molt is confined to June-October. Based primarily on banding studies the suggestions on molt timing in Broad-billed Hummingbird reported by Powers and Wethington (2020) are more consistent with the results of this study, though substantial clarification of preformative, second prebasic, and definitive prebasic molt strategies is provided here. Similar discrepancies between results reported here and those of these previous sources are found in the other seven species. Also contrasting with previous reports, I found that suspended or arrested molts to be rare in these eight species of hummingbirds (< 0.1%), and I also provide new criteria for age determination and its timing, including those related to development of definitive appearance in first-cycle males, molt limits among wing coverts, and molt clines among the remiges (Supplemental Figures S1-S9). No evidence was found for an identifiable second basic plumage in male Lucifer Hummingbirds and little evidence for this in Rivoli’s Hummingbird, contra Pyle (1997).

### Evolution of molt strategies in hummingbirds

I found no evidence for prealternate molts in these eight species of hummingbirds, although such evidence would be better gained from banding studies; prealternate molts may not be as expected in less-migratory or resident hummingbirds (Johnson and Wolfe 2018). Irrespective of this, I believe that the preformative and prebasic molt strategies documented here support the interpretation of Sieburth and Pyle (2018) that the second prebasic molt has been temporally advanced in northern hummingbirds of the United States, as opposed to traditional interpretations that the first complete molt of North American hummingbirds is invariably the preformative molt. In Rivoli’s Hummingbird and Blue-throated Mountain-gem, a partial preformative molt and a complete second prebasic molt averaging earlier in timing than definitive prebasic molts, at about a year of age, is consistent with molt strategies in many other birds, as are complete preformative and prebasic molts during the same temporal period in White-eared Hummingbird. The second prebasic molt in these species peak in August (Figure 2), whereas this molt is here interpreted as peaking progressively earlier in Broad-billed Hummingbird (June), Violet-crowned Hummingbird (May), Berylline Hummingbird (March), Buff-bellied Hummingbird (February), and Lucifer Hummingbird (January), in the last species similar to the timing for the first primary molt in the eight northern species.

Like the northern species (Sieburth and Pyle 2018), Lucifer Hummingbird is highly migratory and undergoes a partial preformative molt of feathers (e.g., those of the gorget in males), that get replaced again during the first molt of primaries in winter and early spring. In order to best preserve homology under the traditional interpretation, the partial-to-incomplete molt of first-cycle Rivoli’s Hummingbirds, Blue-throated Mountain-gems, and Violet-crowned Hummingbirds, and the partial-to-complete molt of first-cycle Broad-billed Hummingbirds, would also be considered auxiliary prefomative molts, which would be novel interpretations for these molts. Rather, I conclude it more parsimonious to interpret the partial-to-complete first-cycle molts that occur primarily in May-December to be preformative molts, as in many other bird species, and that the complete second prebasic molt has evolved along hummingbird lineages to become variably advanced in timing, from August in Rivoli’s Hummingbird and Blue-throated Mountain-gem, to May in Violet-crowned Hummingbird, to January in Lucifer Hummingbird and the other migratory northern species, perhaps in response to the shorter life span of hummingbirds relative to other birds (Sieburth and Pyle 2018).

The eight species of hummingbirds studied here are found in three clades as defined by McGuire et al. (2014), the bee clade (Lucifer Hummingbird), mountain-gem clade (Rivoli’s Hummingbird and Blue-throated Mountain-gem), and emerald clade (remaining five species), with the emeralds being split into four groups as defined by Stiles et al. (2017), including group A (Broad-billed Hummingbird), group B (White-eared Hummingbird), and group D (Violet-crowned, Berylline, and Buff-bellied hummingbirds). Molt strategies in these eight species could be interpreted as having differentiated through synapomorphy during the evolution of these clades and groups, with the bee clade (including the eight northern North American species) sharing more limited preformative molts and second prebasic molts at 6-8 months of age, the mountain-gem clade sharing partial preformative molts and second prebasic molts at about a year of age, and the emerald clade sharing molt strategies that differentiate according to group, with variable preformative molts and second prebasic molts at a year of age (group A), complete preformative molts (group B), or protracted and partial preformative molts followed by second prebasic molts that occur at 7-10 months of age (group D). Partial preformative molts in the more-primitive topaz, hermit, and patagona clades (Zimmer 1950, Hu et al. 2000, Pyle et al. 2015, Johnson and Wolfe 2018) could represent the ancestral state (Sieburth and Pyle 2018).

Within the emerald clade, however, Johnson and Wolfe (2018) indicate that at least one species in group B (genus *Campylopterus*) may have a partial preformative molt and at least three species in group D (now in genera *Chrysuronia, Chionomesa,* and *Hylocharis)* may have complete preformative molts, contrasting with the above-proposed shared molt-strategy partitioning for the species in this study. Molt strategies on many more species of hummingbirds will need to be documented to further test how they have evolved along ancestral Trochilid lineages. Given the plastic nature of molt strategies found by these and other studies on avian molt to date, within genera and even within species (Johnson 1985, Voelker and Rohwer 1998, Rohwer and Irving 2011, Rohwer et al. 2011), I predict that variation in the extent and timing of preformative molts and the timing of prebasic molts in hummingbirds will be shaped more by environmental factors than by synapomorphy.

### Analysis of digital images to study bird molt

As shown by the results of this study, the Macaulay Library and eBird checklists clearly provide an important resource for the investigation of avian molts and plumages, particularly with respect to sequence of remigial replacement, the extent of partial and incomplete molts, the timing of complete molts, and plumage-related criteria for age determination. Certain aspects of molt strategies will still need to be assessed through specimens, in which, for example, age and reproductive status can be confirmed with extent of bill corrugations and information about gonads and other conditions recorded on specimen labels. Data from banding studies, furthermore, can add information on known individuals through recaptures, and I predict that exceptions to some of the information presented here will be found during these studies. Analyses of individual feathers for stable isotopes and connectivity between summer and winter grounds can be undertaken with specimens and captured birds but not with images. Additional drawbacks to scoring molt from images include the quality of some images, making it difficult or impossible to determine precise remegial numbering, the inability to assess both wings to confirm symmetrical molt for many individuals, difficulty in assessing low levels of body-feather molt, and in the case of hummingbirds, the effects that lighting can have on iridescent display feathers as presented in single-plane images. However, these concerns are mitigated by the substantial sample sizes of available images, resulting in adequate data despite the usability of only small proportions of these samples, and, in many cases, the ability to assess multiple images of the same individual in one or more eBird checklists.

Both specimen examination and banding studies take time and effort, as opposed to examination of on-line images, during which large samples can be gathered and analyzed in a short amount of time and with little expense, data are collected without having to be concerned about damaging specimens or the health of a captive bird, and voucher photographs are automatically part of the methodology and can be preserved for later examination or studies on repeatability of results. As shown by the validation study reported on here, banders and field ornithologists of varying experience levels can collect accurate data from images, with precision of data appearing to increase with experience levels of banding and (less so) field experience. I also predict that similar validation studies performed with specimens and banded birds would yield similar levels of accuracy.

I encourage additional research on avian molt though examination of digital images. Here I provide a road map for a subset of such studies; however, many other questions on molt can also be investigated using the Macaulay Library collection. For example, how might timing of molt in these eight species of hummingbirds vary with respect to breeding and wintering locations or in subtropical and tropical breeding subspecies or populations (cf. Wagner 1957, Guallar and Gallés 2017)? How much molt-breeding overlap may occur for birds photographed repeatedly at known nesting sites (e.g., ([ML174305101](https://macaulaylibrary.org/asset/174305101))? How might remigial replacement sequence vary in little-known bird families, and can this be applied to the evolution of molt sequence and of birds? Data from the Macaulay Library image collection can also supplement other data sets to help answer questions related to molt intensity and duration (Rohwer et al. 2009) and to the evolution of preformative molts and formative plumages through phylogenetic comparative or ancestral state reconstruction analysis (cf. Kiat et al. 2019), as have recently been performed based on specimens in other New World bird families such as Cardinalidae (Guallar et al. 2020) and Parulidae (Terrill et al. 2020). To best further such research, finally, I encourage those contributing images to eBird to include birds in molt or in worn plumages, even if they may not be as appealing as, for example, adult males in definitive plumage, of which >50% of hummingbird images I examined referred.

## Acknowledgments

Foremost I thank the thousands of citizen scientists who have contributed images to the Macaulay Library and for agreeing to the license allowing use for research purposes. A total of 174 contributors provided images that are shown or linked in this study. See Figures 2-3, Supplemental Table S2, and Supplemental Figures S1-S9 for contributors to those media. Photos linked in the primary manuscript and for the validation exercise, used by license agreement from the Macaulay Library, were contributed by © Ryan Andrews, Eric Barnes, Cathy Beck, James W. Beck, David Bernstein, Robert Bowling, Paul Budde, Cesar Castillo, Fred Collins, Ed Corey, Joshua Covill, Holly Cox, Jon Curd, Tom Driscoll, Merryl Edelstein, Laura Ellis, Richard Fray, Tony Godfrey, Bradley Hacker, Brien Harvey, John Haynes, Bill Hubick, Scott Jennex, Eric Kallen, Ad Konings, Scott Lewis, Anuar López, Suzie McCann, Michael McCloy, Ken Murphy, Marky Mutchler, Grace Oliver, Arlene Ripley, Dan Scheiman, Mel Senac, John Sterling, James Stull, John Sullivan, Brandon Trentler, Mary Rachel Tucker, Jason Vassallo, Dan Vickers, Nigel Voaden, Rob Worona, Bill Ypsilantis, and Bob Zaremba; see the Supplemental Materials File for additional contributors. I thank Matthew Medler and Brooke Kelley Keeney of the Cornell Lab for help ensuring that information about the Macaulay Library and contributors through eBird were accurately and properly credited, and Blaine Carnes, Jerry Cole, Kenneth Foster, Marcel Gahbauer, Christine Godwin, Santiago Guallar, Marc Illa, Erik Johnson, Danielle Kaschube, Brooke Kelley Keeney, Pricilla Lai, Shannon M. Mendia, Jim Saracco, Dessi Sieburth, Al Sherkow, Natália Pérez Ruiz, Ron Taylor, and Keegan Tranquillo for participating in the validation exercise. Blaine Carnes and Santiago Guallar have read a draft of the manuscript and provided constructive criticism. This is contribution # 678 of The Institute for Bird Populations.

