## Supplemental Materials File for "Examination of digital images from Macaulay Library to determine avian molt strategies: A case study on molts and plumages in eight species of North American hummingbirds"

Contents

Table S1. Sample sizes for images and molt and plumage states, by month, for each of eight species of hummingbird that breed in the southwestern United States.

Table S2. Earliest date, latest date, links to voucher images, and days duration recorded for each plumage and molt state for each of eight species of hummingbird that breed in the southwestern United States.

Figure S1. Images exemplifying molts and plumages in Rivoli's Hummingbird.

Figure S2. Images exemplifying molts and plumages in Blue-throated Mountain-gem.

Figure S3. Images exemplifying molts and plumages in Lucifer Hummingbird.

Figure S4. Images exemplifying molts and plumages in Broad-billed Hummingbird.

Figure S5. Images exemplifying molts and plumages in White-eared Hummingbird.

Figure S6. Images exemplifying molts and plumages in Violet-crowned Hummingbird.

Figure S7. Images exemplifying molts and plumages in Berylline Hummingbird.

Figure S8. Images exemplifying molts and plumages in Buff-bellied Hummingbird.

Figure S9. Images of southwestern hummingbirds in formative male plumages.

Validation exercise to assess whether or not field ornithologists can accurately collect data on avian molt from images

Literature cited

Acknowledgements

**Supplementary Table S1. Sample sizes for images and molt and plumage states, by month, for each of eight species of hummingbird that breed in the southwestern United States.**

| | JAN | FEB | MAR | APR | MAY | JUN | JUL | AUG | SEP | OCT | NOV | DEC | TOTAL |
|---|---|---|---|---|---|---|---|---|---|---|---|---|---|
| **Rivoli's Hummingbird (*Eugenes fulgens*)** | | | | | | | | | | | | | |
| *Number of Images* | *201* | *151* | *474* | *527* | *542* | *425* | *605* | *569* | *226* | *171* | *170* | *185* | *4246* |
| Juvenile Plumage | | | | 5 | 6 | 8 | 9 | 5 | 2 | 1 | | | 36 |
| Preformative Molt | 4 | 2 | | | 3 | 6 | 12 | 9 | 11 | 9 | 12 | 9 | 77 |
| Formative Plumage | 2 | 3 | 10 | 18 | 26 | 21 | 13 | 5 | | | | 1 | 99 |
| Second Prebasic Molt | | | | | | 3 | 6 | 9 | 3 | | | | 21 |
| Definitive Basic Plumage | 13 | 9 | 21 | 34 | 42 | 27 | 34 | 24 | 5 | 3 | 12 | 14 | 238 |
| Definitive Prebasic Molt | | | | | | | 3 | 28 | 23 | 7 | 3 | | 64 |
| ***Total Individuals*** | ***19*** | ***14*** | ***31*** | ***57*** | ***77*** | ***65*** | ***77*** | ***80*** | ***44*** | ***20*** | ***27*** | ***24*** | ***535*** |
| **Blue-throated Mountain-gem (*Lamphornis clemenciae*)** | | | | | | | | | | | | | |
| *Number of Images* | *317* | *123* | *108* | *222* | *309* | *231* | *343* | *446* | *111* | *105* | *137* | *91* | *2543* |
| Juvenile Plumage | | | | | 3 | 4 | 5 | 2 | | | | | 14 |
| Preformative Molt | | | | | 1 | 4 | 12 | 32 | 20 | 15 | 7 | 1 | 92 |
| Formative Plumage | 3 | 4 | 7 | 12 | 10 | 11 | 14 | 11 | | | 1 | 4 | 77 |
| Second Prebasic Molt | | | | | | | 1 | 19 | 4 | | | | 24 |
| Definitive Basic Plumage | 8 | 9 | 7 | 19 | 21 | 16 | 29 | 37 | 14 | 3 | 9 | 4 | 176 |
| Definitive Prebasic Molt | | | | | | | | 14 | 16 | 5 | 1 | | 36 |
| ***Total Individuals*** | ***11*** | ***13*** | ***14*** | ***31*** | ***35*** | ***35*** | ***61*** | ***115*** | ***54*** | ***23*** | ***18*** | ***9*** | ***419*** |
| **Lucifer Hummingbird (*Calothrax lucifer*)** | | | | | | | | | | | | | |
| *Number of Images* | *12* | *4* | *124* | *517* | *531* | *333* | *528* | *845* | *438* | *132* | *20* | *10* | *3494* |
| Juvenile Plumage | | | | 14 | 19 | 19 | 25 | 17 | 9 | 4 | | | 107 |
| Preformative Molt | | | | | | 2 | 10 | 57 | 57 | 24 | 1 | | 151 |
| Second Prebasic Molt | 3 | | 1 | | | | | | 1 | | 2 | 3 | 10 |
| Definitive Basic Plumage | | 1 | 20 | 82 | 102 | 70 | 92 | 117 | 39 | 7 | 1 | | 531 |
| Definitive Prebasic Molt | 4 | | | | | | 3 | 58 | 55 | 7 | 4 | 1 | 132 |
| ***Total Individuals*** | ***7*** | ***1*** | ***21*** | ***96*** | ***121*** | ***91*** | ***130*** | ***249*** | ***161*** | ***42*** | ***8*** | ***4*** | ***931*** |

**Supplementary Table S1 (cont.). Sample sizes for images and molt and plumage states, by month, for each of eight species of hummingbird that breed in the southwestern United States.**

| | JAN | FEB | MAR | APR | MAY | JUN | JUL | AUG | SEP | OCT | NOV | DEC | TOTAL |
|---|---|---|---|---|---|---|---|---|---|---|---|---|---|
| **Broad-billed Hummingbird (*Cynanthus latirostris*)** | | | | | | | | | | | | | |
| *Number of Images* | *631* | *424* | *1160* | *1097* | *1028* | *642* | *1184* | *959* | *482* | *473* | *440* | *322* | *8842* |
| Juvenile Plumage | | | 2 | 8 | 16 | 20 | 20 | 11 | 2 | | | | 79 |
| Preformative Molt | 3 | 3 | 0 | 0 | 2 | 10 | 117 | 130 | 38 | 27 | 11 | 7 | 348 |
| Formative Plumage | 24 | 13 | 39 | 69 | 79 | 32 | 12 | 1 | | | 3 | 11 | 283 |
| Second Prebasic Molt | | | | | 1 | 27 | 124 | 99 | 5 | 1 | | | 257 |
| Definitive Basic Plumage | 36 | 39 | 117 | 150 | 137 | 71 | 182 | 60 | 37 | 34 | 36 | 20 | 919 |
| Definitive Prebasic Molt | | | | | | 10 | 184 | 285 | 44 | 4 | | | 527 |
| ***Total Individuals*** | ***63*** | ***55*** | ***158*** | ***227*** | ***235*** | ***170*** | ***639*** | ***586*** | ***126*** | ***66*** | ***50*** | ***38*** | ***2413*** |
| **White-eared Hummingbird (*Basilinna leucotis*)** | | | | | | | | | | | | | |
| *Number of Images* | *40* | *44* | *33* | *20* | *126* | *236* | *259* | *273* | *40* | *16* | *17* | *33* | *1137* |
| Juvenile Plumage | | | 4 | 1 | 1 | 4 | 1 | 3 | 1 | | | | 15 |
| Preformative Molt | | | | | 2 | 8 | 14 | 14 | 4 | 0 | | | 42 |
| Formative/Basic Plumage | 19 | 17 | 12 | 9 | 22 | 25 | 12 | 18 | 9 | 6 | 9 | 12 | 170 |
| Second/Definitive Prebasic Molt | | | | | | 3 | 23 | 20 | 6 | 1 | | | 53 |
| ***Total Individuals*** | ***19*** | ***17*** | ***16*** | ***10*** | ***25*** | ***40*** | ***50*** | ***55*** | ***20*** | ***7*** | ***9*** | ***12*** | ***280*** |
| **Violet-crowned Hummingbird (*Leucolia violiceps*)** | | | | | | | | | | | | | |
| *Number of Images* | *198* | *171* | *354* | *336* | *275* | *128* | *522* | *418* | *126* | *68* | *71* | *170* | *2837* |
| Juvenile Plumage | | | | | 1 | 4 | 8 | 5 | 0 | 0 | 0 | 0 | 18 |
| Preformative Molt | 2 | | | | | 11 | 35 | 36 | 24 | 12 | 4 | 2 | 126 |
| Formative Plumage | 6 | 8 | 24 | 20 | 3 | | | | 3 | 7 | 6 | 11 | 88 |
| Second Prebasic Molt | | | | 18 | 26 | 7 | | | | | | | 51 |
| Definitive Basic Plumage | 7 | 10 | 18 | 27 | 19 | 37 | 67 | 48 | 15 | 6 | 6 | 2 | 262 |
| Definitive Prebasic Molt | | | | | 3 | 6 | 18 | 21 | 6 | 2 | | | 56 |
| ***Total Individuals*** | ***15*** | ***18*** | ***42*** | ***65*** | ***52*** | ***65*** | ***128*** | ***110*** | ***48*** | ***27*** | ***16*** | ***15*** | ***601*** |

**Supplementary Table S1 (cont.). Sample sizes for images and molt and plumage states, by month, for each of eight species of hummingbird that breed in the southwestern United States.**

| | JAN | FEB | MAR | APR | MAY | JUN | JUL | AUG | SEP | OCT | NOV | DEC | TOTAL |
|---|---|---|---|---|---|---|---|---|---|---|---|---|---|
| **Berylline Hummingbird (*Saucerottia beryllina*)** | | | | | | | | | | | | | |
| *Number of Images* | *57* | *43* | *59* | *79* | *339* | *349* | *134* | *304* | *108* | *33* | *48* | *54* | *1607* |
| Juvenile Plumage | | | | | | 1 | | 4 | 1 | | 1 | | 7 |
| Preformative Molt | 2 | | | | | | | 2 | 3 | 6 | 8 | 9 | 30 |
| Formative Plumage | 5 | 2 | 1 | | | | | | | | 1 | 10 | 19 |
| Second Prebasic Molt | 3 | 3 | 10 | 5 | 1 | 1 | | | | | | 1 | 24 |
| Definitive Basic Plumage | 5 | 6 | 11 | 13 | 23 | 31 | 37 | 35 | 19 | 14 | 8 | 4 | 206 |
| Definitive Prebasic Molt | 7 | 1 | 1 | | | | 1 | 2 | 2 | 5 | 15 | 18 | 52 |
| ***Total Individuals*** | ***22*** | ***12*** | ***23*** | ***18*** | ***24*** | ***33*** | ***38*** | ***43*** | ***25*** | ***25*** | ***33*** | ***42*** | ***338*** |
| **Buff-bellied Hummingbird (*Amazilia yucatanensis*)** | | | | | | | | | | | | | |
| *Number of Images* | *563* | *425* | *305* | *365* | *120* | *47* | *42* | *33* | *113* | *142* | *459* | *261* | *2875* |
| Juvenile Plumage | | 3 | 7 | 5 | | | 1 | | | | | | 16 |
| Preformative Molt | | 2 | 4 | 4 | 7 | 4 | 7 | 5 | 8 | 12 | 18 | 3 | 74 |
| Formative Plumage | 40 | 23 | 16 | 20 | 4 | | 1 | 2 | 3 | 7 | 23 | 30 | 169 |
| Second Prebasic Molt | 6 | 8 | 2 | | 1 | | | | | | 2 | 3 | 22 |
| Definitive Basic Plumage | 77 | 81 | 62 | 79 | 28 | 14 | 12 | 8 | 14 | 12 | 58 | 48 | 493 |
| Definitive Prebasic Molt | 0 | 1 | 1 | 2 | | | 1 | 1 | 10 | 13 | 19 | 6 | 54 |
| ***Total Individuals*** | ***123*** | ***118*** | ***92*** | ***110*** | ***40*** | ***18*** | ***22*** | ***16*** | ***35*** | ***44*** | ***120*** | ***90*** | ***828*** |
| **Total Individuals (all species)** | **279** | **248** | **397** | **614** | **609** | **517** | **1145** | **1254** | **513** | **254** | **281** | **234** | **6345** |

**Table S2. Earliest date, latest date, links to voucher images, and days duration recorded for each plumage and molt state for each of eight species of hummingbird that breed in the southwestern United States.**

| | Earliest Date | Image Link | Latest Date | Image Link | No. days[1] |
|---|---|---|---|---|---|
| **Rivoli's Hummingbird (*Eugenes fulgens*)** | | | | | |
| Juvenile Plumage | 5 Apr (2017) | ML53539161 | 6 Oct (2017) | ML71217491 | 184 |
| Preformative Molt | 19 May (2009) | ML35507671 | 25 Feb (2018) | ML87527351 | 282 |
| Formative Plumage | 27 Dec (2018) | ML131888891 | 27 Aug (2017) | ML67175761 | 130 |
| Second Prebasic Molt | 8 Jun (2020) | ML242091451 | 15 Sep (2017) | ML69429691 | 99 |
| Definitive Basic Plumage | 7 Sep (2018) | ML115560961 | 1 Sep (2018) | ML114745281 | 359 |
| Definitive Prebasic Molt | 16 Jul (2020) | ML250033061 | 22 Nov (2019) | ML189631681 | 129 |
| **Blue-throated Mountain-gem (*Lamphornis clemenciae*)[2]** | | | | | |
| Juvenile Plumage | 12 May (2018) | ML 104413711 | 23 Aug (2019) | ML174516651 | 103 |
| Preformative Molt | 31 May (2008) | ML116833491 | 31 Dec (2018) | ML132225371 | 214 |
| Formative Plumage | 23 Nov (2018) | ML125145541 | 26 Aug (2016) | S31263149 | 276 |
| Second Prebasic Molt | 23 Jul (2016) | ML91903061 | 24 Sep (2017) | ML70364721 | 63 |
| Definitive Basic Plumage | 30 Sep (2017) | ML70410371 | 18 Sep (2014) | ML61572141 | 353 |
| Definitive Prebasic Molt | 2 Aug (2019) | ML177245281 | 4 Nov (2018) | ML125171011 | 94 |
| **Lucifer Hummingbird (*Calothrax lucifer*)[3]** | | | | | |
| Juvenile Plumage | 14 Apr (2016) | ML27082041 | 14 Oct (2016) | ML37413671 | 183 |
| Preformative Molt | 5 Jun (2020) | S70103132 | 4 Nov (2017) | ML74135881 | 152 |
| Second Prebasic Molt | 8 Sep (2017) | ML68546921 | 2 Mar (2018) | ML88246211 | 176 |
| Definitive Basic Plumage | 28 Feb (2019) | ML143090201 | 15 Nov (2015) | ML21063081 | 260 |
| Definitive Prebasic Molt | 3 Jul (2020) | ML247351211 | 20 Jan (2020) | ML200761501 | 201 |
| **Broad-billed Hummingbird (*Cynanthus latirostris*)** | | | | | |
| Juvenile Plumage | 16 Mar (2020) | ML215925901 | 6 Sep (2016) | ML 34443551 | 143 |
| Preformative Molt | 26 Jun (2020) | ML245827291 | 19 Feb (2018) | ML86692301 | 238 |
| Formative Plumage | 14 Nov (2016) | ML39926711 | 1 Aug (2019) | ML197319181 | 260 |
| Second Prebasic Molt | 21 May (2018) | ML101361191 | 2 Oct (2018) | ML 120044251 | 134 |
| Definitive Basic Plumage | 27 Jul (2020) | ML252081611 | 24 Aug (2005) | ML87864221 | 393 |
| Definitive Prebasic Molt | 7 Jun (2019) | ML167008081 | 10 Oct (2019) | ML183630481 | 156 |
| **White-eared Hummingbird (*Basilinna leucotis*)[4]** | | | | | |
| Juvenile Plumage | 9 Mar (2020) | ML220372891 | 11 Sep (2019) | S59696591 | 186 |
| Preformative Molt | 28 May (2020) | ML239528951 | 24 Sep (2015) | S30924176 | 148 |
| Formative/Basic Plumage | 10 Aug (2008) | ML136971191 | 19 Aug (2019) | ML173276031 | 374 |
| Second/Definitive Prebasic Molt | 3 Jun (2013) | ML222523661 | 6 Oct (2019) | ML185831481 | 125 |
| **Violet-crowned Hummingbird (*Leucolia violiceps*)** | | | | | |
| Juvenile Plumage | 30 May (2014) | ML 34273361 | 16 Aug (2018) | ML112325371 | 78 |
| Preformative Molt | 20 Jun (2020) | ML244744561 | 24 Jan (2020) | ML202657101 | 218 |
| Formative Plumage | 11 Sep (2019) | ML213615071 | 29 May (2009) | ML111468491 | 260 |
| Second Prebasic Molt | 9 Apr (2009) | ML 191090781 | 13 Jun (2020) | ML243892441 | 65 |
| Definitive Basic Plumage | 11 June (2018) | ML131533621 | 17 Sep (2016) | ML62598551 | 267 |
| Definitive Prebasic Molt | 3 May (2010) | ML245135201 | 28 Oct (2019) | ML185533471 | 178 |

**Table S2 (cont.). Earliest date, latest date, links to voucher images, and days duration recorded for each plumage and molt state for each of eight species of hummingbird that breed in the southwestern United States.**

| | Earliest Date | Image Link | Latest Date | Image Link | No. days[1] |
|---|---|---|---|---|---|
| **Berylline Hummingbird (*Saucerottia beryllina*)** | | | | | |
| Juvenile Plumage | 27 Jun (2020) | ML246032741 | 18 Nov (2018) | ML124156711 | 144 |
| Preformative Molt | 18 Aug (2019) | ML173051291 | 21 Jan (2018) | ML82872751 | 156 |
| Formative Plumage | 20 Nov (2017) | ML75733781 | 16 Mar (2010) | ML24493361 | 116 |
| Second Prebasic Molt | 12 Dec (2019) | ML192834001 | 9 Jun (2020) | ML242471461 | 186 |
| Definitive Basic Plumage | 22 Jan (2020) | ML210693961 | 22 Dec (2015) | ML22259701 | 334 |
| Definitive Prebasic Molt | 10 Jul (2017) | ML63020951 | 10 Mar (2015) | ML46645931 | 243 |
| **Buff-bellied Hummingbird (*Amazilia yucatanensis*)** | | | | | |
| Juvenile Plumage | 2 Feb (2002) | ML23055171 | 14 Jul (2010) | ML 126355861 | 162 |
| Preformative Molt | 11 Feb (2017) | ML48095981 | 17 Dec (2017) | ML78493091 | 309 |
| Formative Plumage | 20 Jul (2016) | S30779253 | 19 May (2020) | S69336911 | 333 |
| Second Prebasic Molt | 4 Nov (2016) | ML39289491 | 11 May (2017) | ML57657391 | 188 |
| Definitive Basic Plumage | 10 Sep (2006) | ML163757271 | 26 Nov (2003) | ML34434691 | 442 |
| Definitive Prebasic Molt | 29 Jul (2018) | ML109577131 | 26 Apr (2016) | ML 44139031 | 271 |

[1] Days duration from earliest to latest dates in table (non-leap-year calendar). Note that values can be > year for conditions showing substantial individual variation in timing (e.g., for definitive basic plumage).

[2] Post-fledging male Blue-throated Mountain-gems with blue iridescent feathers in the throat were categorized as in formative rather juvenile plumage (see Supplemental Figure S9).

[3] No line is presented for formative Lucifer Hummingbirds due to lack of sample sizes.

[4] No lines are presented for White-eared Hummingbirds for formative plumage or undergoing the second prebasic molt as these states can not be identified in photographs (see text and Supplemental Figure S5).

**Figures S1-S8. Images of southwestern hummingbirds exemplifying different molts and plumages and useful for age determination.** Species are: Figure S1 - Rivoli's Hummingbird (*Eugenes fulgens*), Figure S2 - Blue-throated Mountain-gem (*Lamphornis clemenciae*), Figure S3 - Lucifer Hummingbird (*Calothrax lucifer*), Figure S4 - Broad-billed Hummingbird (*Cynanthus latirostris*), Figure S5 - White-eared Hummingbird (*Basilinna leucotis*), Figure S6 - Violet-crowned Hummingbird (*Leucolia violiceps*), Figure S7 - Berylline Hummingbird (*Saucerottia beryllina*), and Figure S8 - Buff-bellied Hummingbird (*Amazilia yucatanensis*).

For most species six images are shown, exemplifying juvenile plumage, preformative molt, formative plumage, second prebasic molt, definitive basic plumage, and definitive prebasic molt. Broad-billed Hummingbird shows more variation to the extent of the preformative molt, hence 12 images are shown in Figure S4. In White-eared Hummingbird plumages cannot be identified after the complete preformative molt so additional images of birds undergoing this molt are shown in Figure S5. Dates of images shown are typical of the timing for each of these molts and plumages.

Brief notes on age-determination criteria are noted under each image. Males are shown for the most part; however criteria related to wing feathers apply to ageing females as well. New criteria emphasized include differences between juvenile and basic wing feathers, molt limits among greater coverts, contrasts between newer secondaries and older primaries (s1-p1 contrast), and in some cases molt clines among primaries. Juvenile outer primaries and secondaries tend to be much browner and more worn than replaced basic primaries of the same age, which are duskier and often more lustrous. Many hummingbirds that undergo partial molts replace at least some upperwing lesser coverts and retain at least some greater coverts, resulting in molt limits within these tracts that are useful for ageing; coverts generally appear to be replaced in a distal-caudal direction, as in passerines (see Figure X in Pyle 1997). In birds undergoing protracted molt, an s1-p1 contrast results from the secondaries being replaced well after the inner primaries, in hummingbirds not until p6-p7 are being replaced, by which time inner primaires can be 2-3 months old and show more wear. Likewise, molt clines can also be visible among primaries after protracted molts, with p9, the last feather replaced in hummingbirds, being fresher and appearing slightly darker than both p8 and p10. Birds with juvenile remiges, in contrast, have uniform feathers which were all grown at the same time and show no contrasts or clines due to age; if anything, secondaries are paler and browner than primaries after several months post-fledging due to increased solar exposure. These differences can help distinguish formative from definitive basic plumages in hummingbirds, especially in females, which often lack age-related differences in appearance of body feathering (cf. Supplemental Figure S9).

Links to Macaulay Library specimen pages (ML followed by 8 or 9 numerals) for each image are provided and can be used to view enlarged version of each image, along with the photographer and more detail on location and other aspects of the record.

### Figure S1. Images exemplifying molts and plumages in Rivoli's Hummingbird.

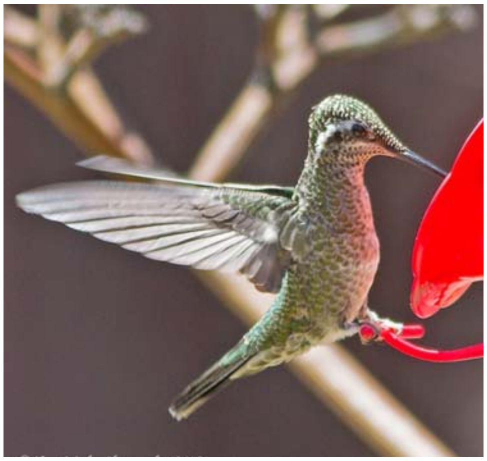

Juvenile female. Note fresh scaly plumage (juvenile males have darker scalloping). 22 May 2015, Arizona, ML193444241.

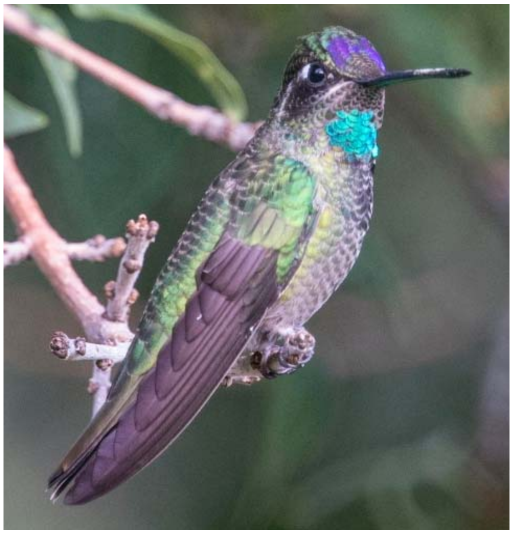

Male undergoing preformative molt. Note incoming gorget feathers, fresh remiges and duller greater coverts. 4 Aug 2020, Arizona, ML171128561.

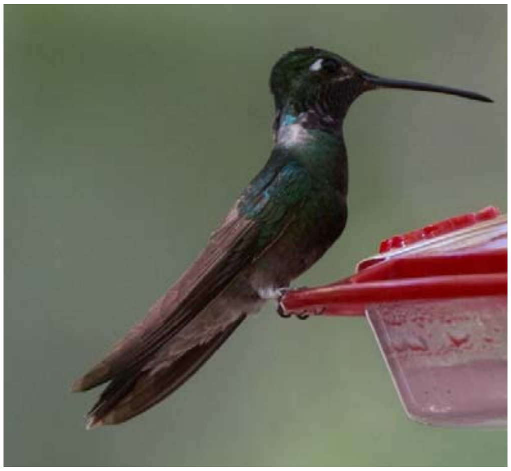

Formative male. Note retained brown juvenile primaries and secondaries. See also Figure 2. 19 May 2013, Arizona, ML42934871.

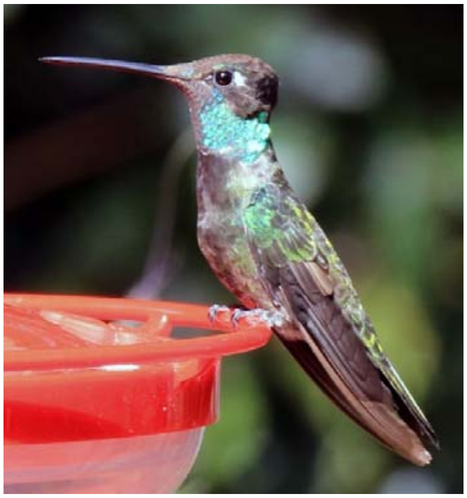

Male undergoing second prebasic molt. Note unmolted juvenile p9-p10 and s3-s4; p8, s2, and s5 are growing. 16 Sep 2017, Arizona, ML69483461.

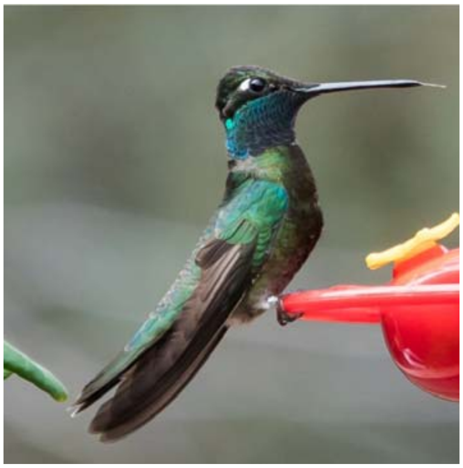

Definitive basic male. Note contrast between newer secondaries and older primaries. 8 Mar 2018, Arizona, ML90525171.

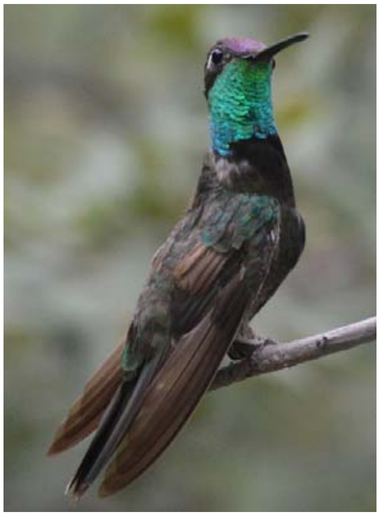

Male undergoing definitive prebasic molt. Note unmolted remiges are duskier than retained juvenile feathers during second prebasic molt. 22 Aug 2015, Arizona, ML128951341.

**Figure S2. Images exemplifying molts and plumages in Blue-throated Mountain-gem.**

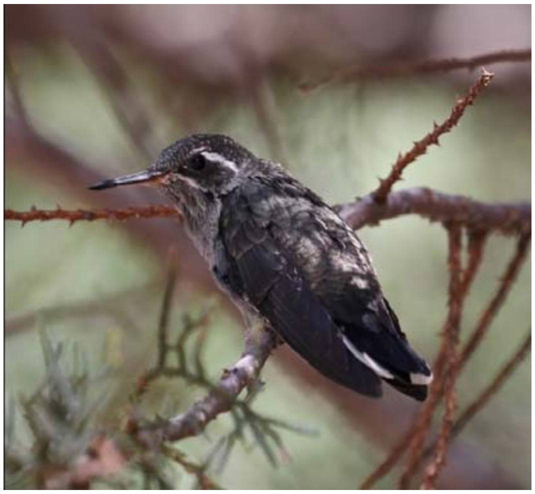

Juvenile. None of 14 juveniles examined showed blue in throat, indicating these may be formative feathers. 1 Aug 2016, Arizona, ML33117161.

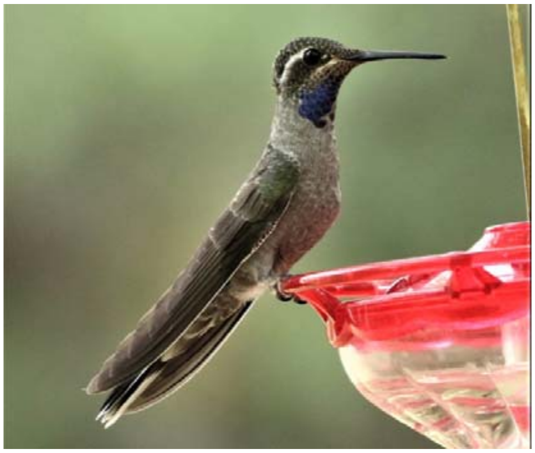

Male undergoing preformative molt. Note incomplete gorget, fresh wings, and duller greater coverts. 2 Sep 2017, Arizona, ML71224011.

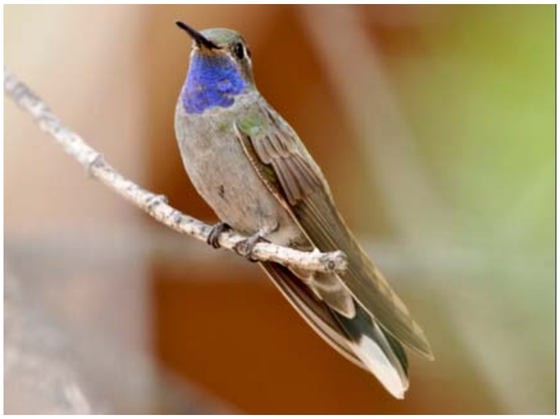

Formative male. Note incomplete gorget, molt limit in wing coverts, and retained brown juvenile primaries and secondaries. See also Figure 2. 14 Jul 2018, Arizona, ML195618151.

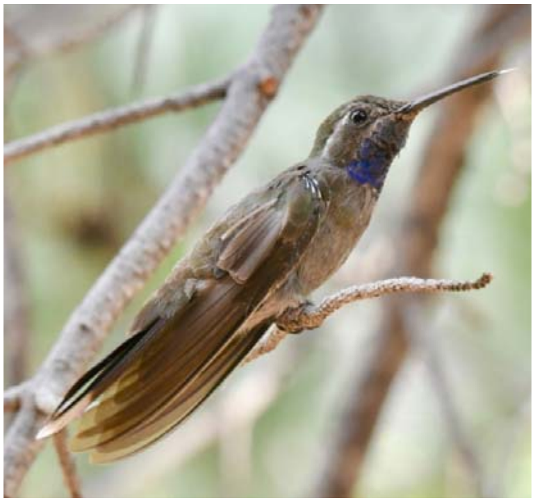

Male commencing second prebasic molt, with p1-3 growing. Note incomplete gorget and worn juvenile outer primaries, secondaries, and remaining greater covert. 13 Aug 2019, Arizona, ML172907241.

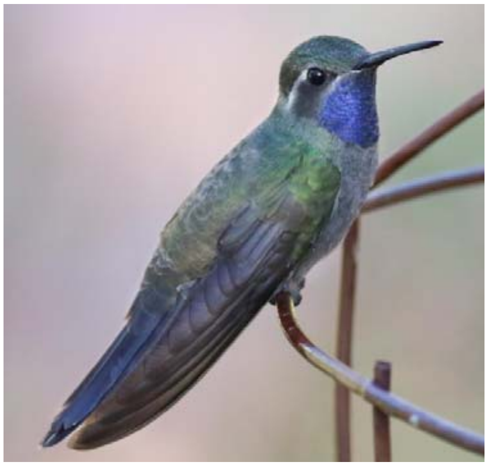

Definitive basic male. Note full gorget and lustrous dark remiges. 4 Jun 2016, Arizona, ML29989261.

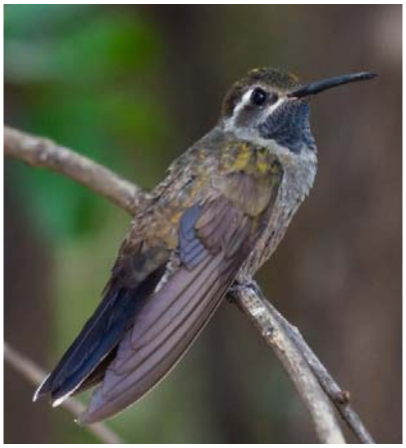

Male undergoing definitive prebasic molt. Note relatively dark and unworn outer primaires and secondaries. 9 Aug 2018, Arizona, ML116501471.

**Figure S3. Images exemplifying molts and plumages in Lucifer Hummingbird.**

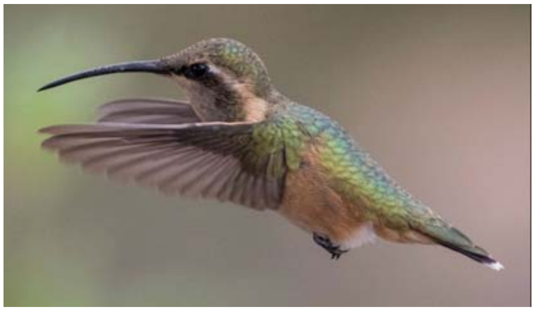

Juvenile. Note fresh scaly plumage and buff underparts. 7 Oct 2015, Arizona, ML20623111.

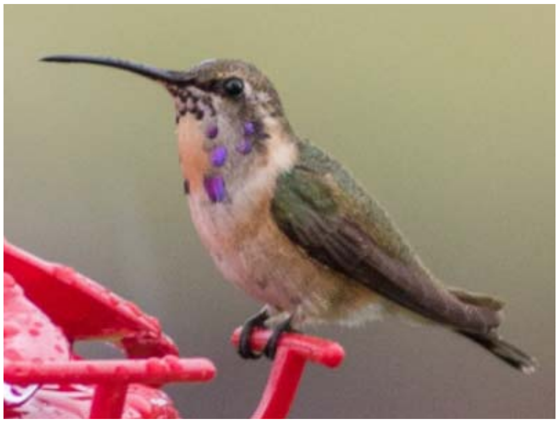

Male undergoing preformative molt. Note uniformly brownish remiges, mixed juvenile and formative (brighter green) back feathers, and formative gorget feathers. Beware occasional definitive basic females can also show purple gorget feathers. 13 Aug 2016, Texas, ML36340281.

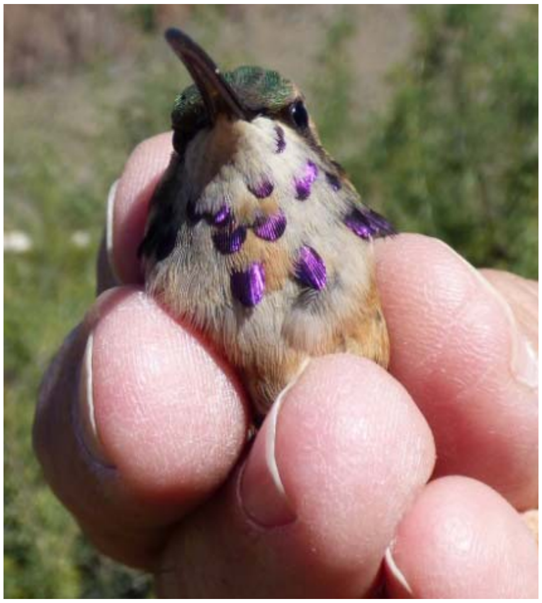

Male undergoing preformative molt or in formative plumage. Crown feathers appear primarily formative, along with a number of gorget feathers. See also Figure 2. 3 Nov 2015, Texas, ML20744131.

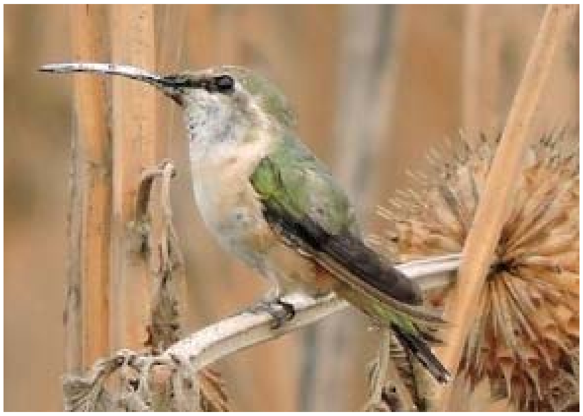

Female undergoing second prebasic molt. Note unmolted, worn brown juvenile p9-p10 and s4; p8 is growing. 2 Mar 2018, Distrito Federal, Mexico, ML88246211.

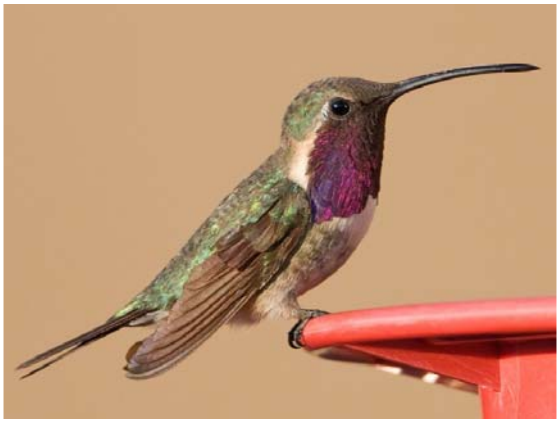

Definitive basic male. Note full gorget, contrast between newer secondaries and older primaries, and molt cline within primaires, p9 appearing newest, a sign of a complete prebasic molt. 26 Aug 2009, Arizona, ML174774571.

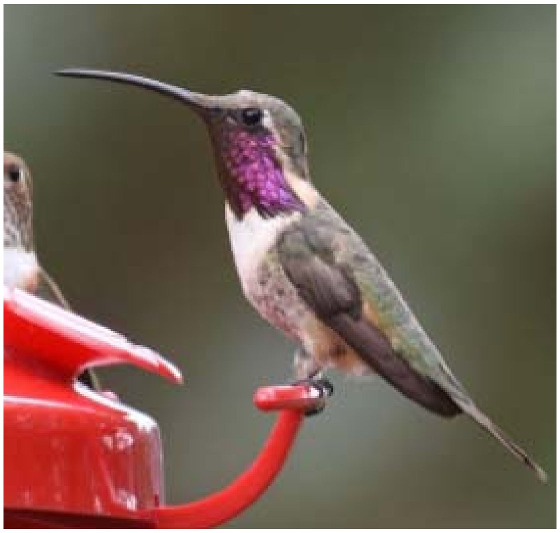

Male undergoing definitive prebasic molt. Note full gorget and dusky basic primaries and secondaries. 7 Sep 2019, Texas, ML176897521.

**Figure S4. Images exemplifying molts and plumages in Broad-billed Hummingbird.**

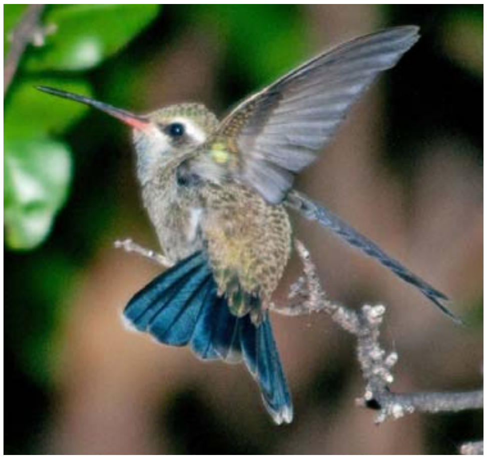

Juvenile male. Note fresh scaly plumage. Males can have a blue wash to juvenile throat feathers but these are not structured like formative gorget feathers.12 May 2016, Arizona, ML35544921.

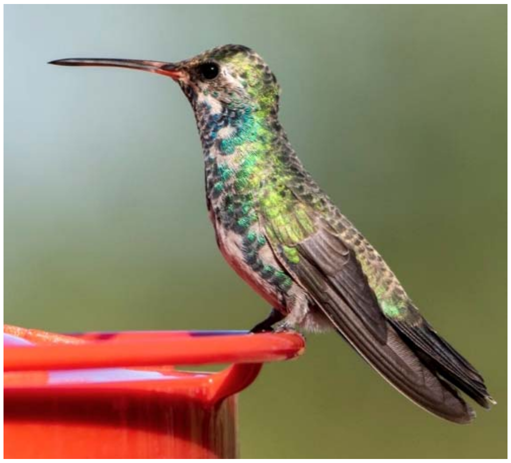

Male undergoing incomplete or complete preformative molt, with p1-p6 and central rectrices being replaced along with body feathering. 27 Aug 2019, Arizona, ML174429551.

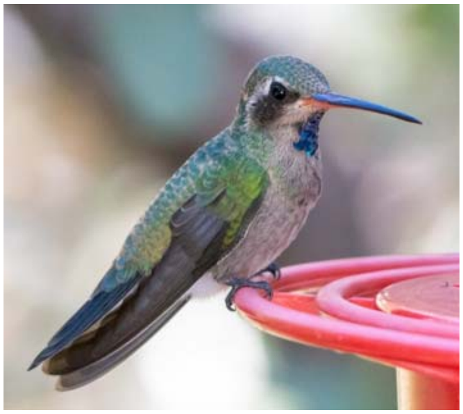

Male commencing preformative molt. Note mixed juvenile and  formative upperpart feathers, incoming gorget feathers on throat, and fresh remiges. 30 Jun 2020, Arizona, ML246780721.

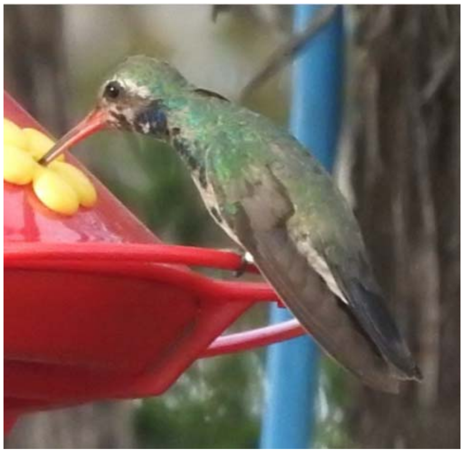

Male undergoing partial preformative molt, replacing body feathers but retaining juvenile flight feathers. 17 Oct 2019, Colorado, ML182857441.

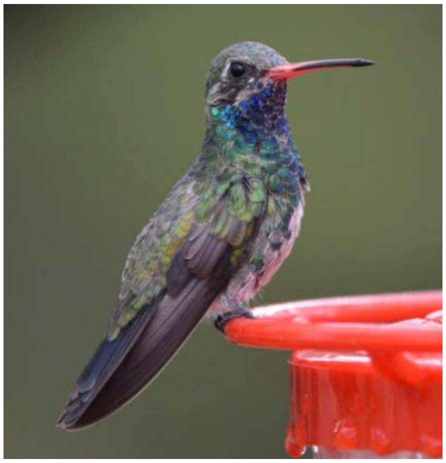

Male undergoing incomplete or complete preformative molt, with p1-p4 and central rectrices being replaced along with body feathering. See also Figure 3. 15 Aug 2019, Arizona, ML172702421.

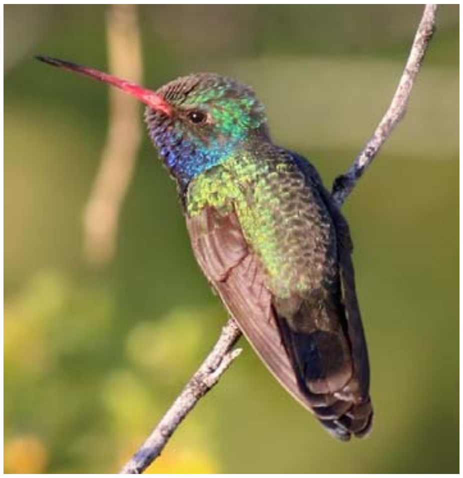

Formative male following partial preformative molt. Note retained juvenile flight feathers and molt limits among upperwing coverts. See also Figure 2. 22 Apr 2014, Arizona, ML226047701.

**Figure S4 (cont.). Molts and plumages in Broad-billed Hummingbird.**

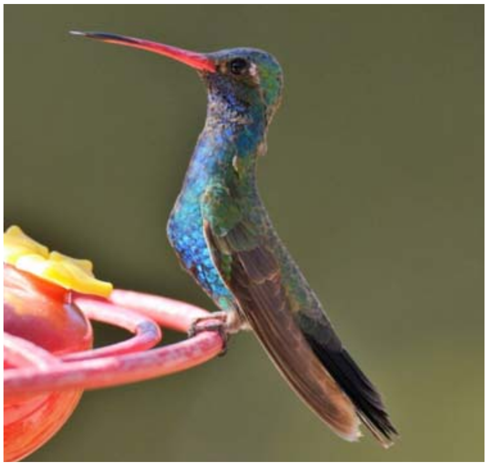

Formative male. Note retained juvenile remiges but replaced formative upperwing coverts. 21 Jun 2020, Arizona, ML246664461.

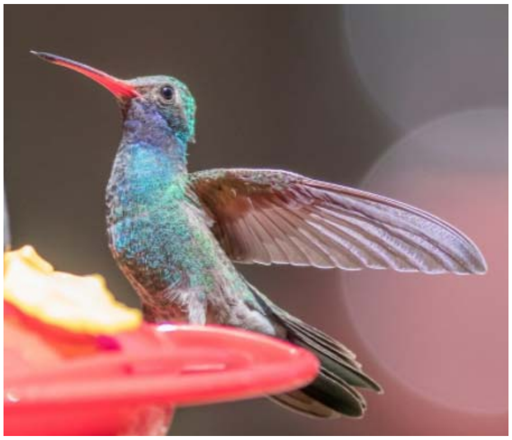

Formative male following complete preformative molt. All flight feathers have been replaced but body feathering appears predefinitive. 6 May 2020, Arizona, ML233606291.

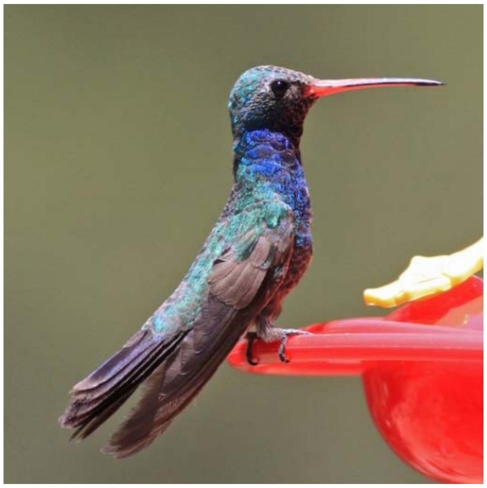

Male undergoing second prebasic molt. Note unmolted, worn and brown, juvenile primaries, secondaries, and central greater covert. 10 Jul 2019, Arizona, ML168722681.

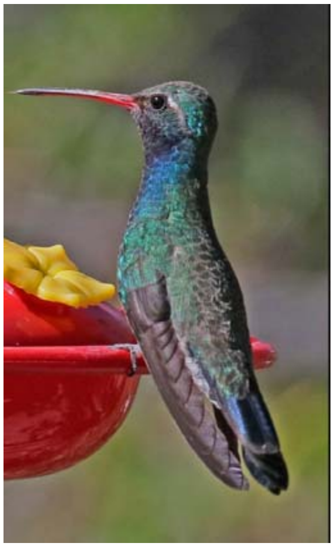

Definitive basic or formative male. Note uniform basic upperwing coverts, darker secondaries than inner primaries, and darker p9 as part of molt cline. Formative males following complete molts may be indistinguishable from definitive basic males. 24 Mar 2019, Arizona, ML148402901.

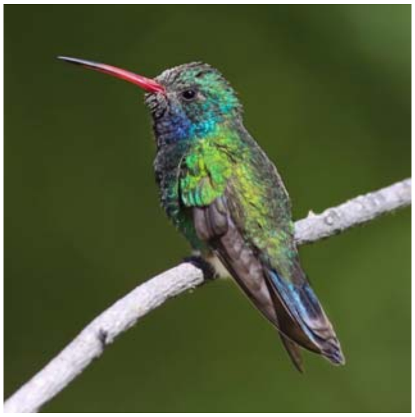

Definitive basic or formative male. Note uniform secondary coverts and darker secondaries than primaries. 23 May 2016, Arizona, ML126747531.

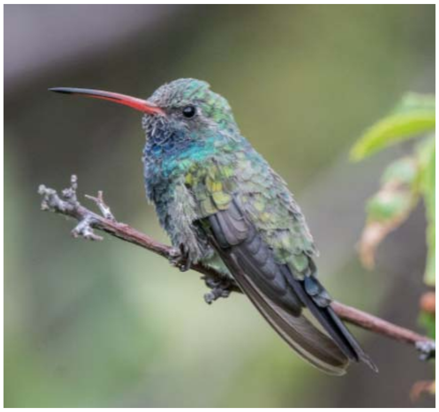

Male undergoing definitive or second prebasic molt. Note non-juvenile feathers; it is possible that body feathering is predefinitive indicating a second prebasic molt. 9 Aug 2019, Arizona, ML171934321.

### Figure S5. Images exemplifying molts and plumages in White-eared Hummingbird.

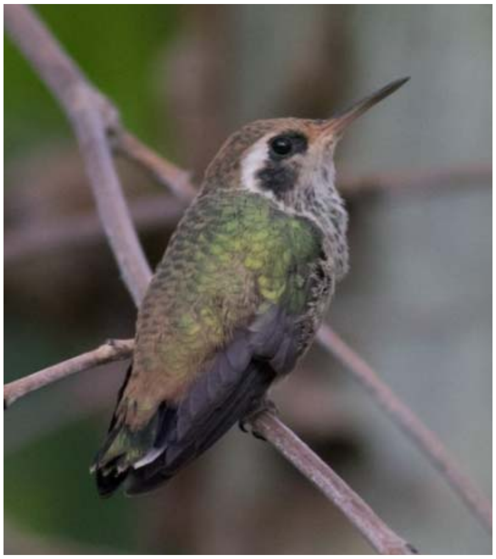

Juvenile. Note fresh scaly plumage and dull bill. 21 Aug 2019, Arizona, ML173592171.

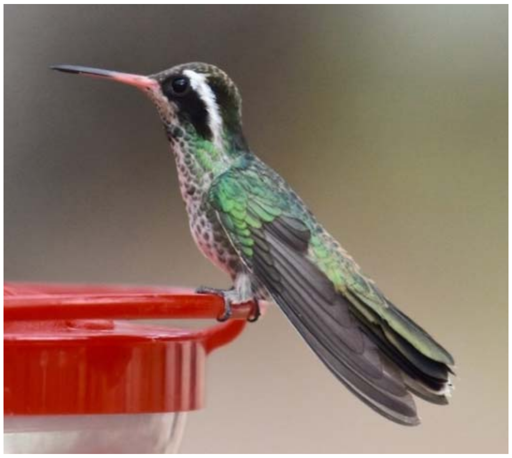

Male undergoing preformative molt. Note concurrence of body feather and primary molt (p1-p3 dropped) as part of complete molt. 13 Jun 2019, Arizona, ML168126461.

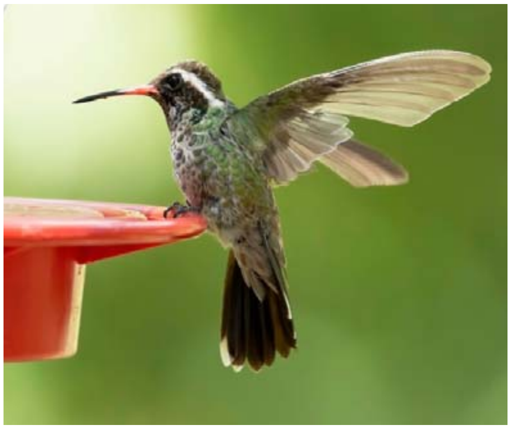

Male undergoing preformative molt. P1-p5 are molting at the same time that body feathering is molting. 9 Aug 2020, Arizona, ML255139331.

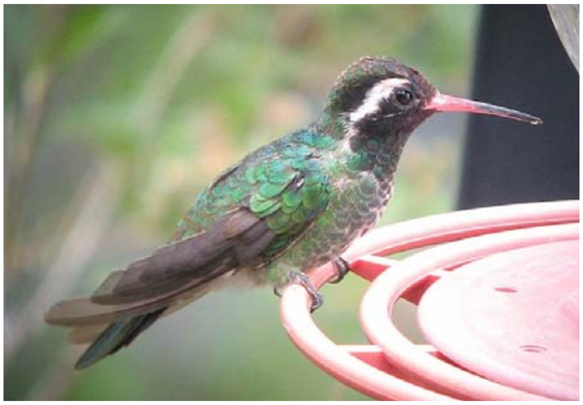

Male completing preformative molt. The juvenile p9 and s4 remain while body-feather molt into a definitive male appearance has almost completed. 2 Aug 2009, Arizona, ML112506621.

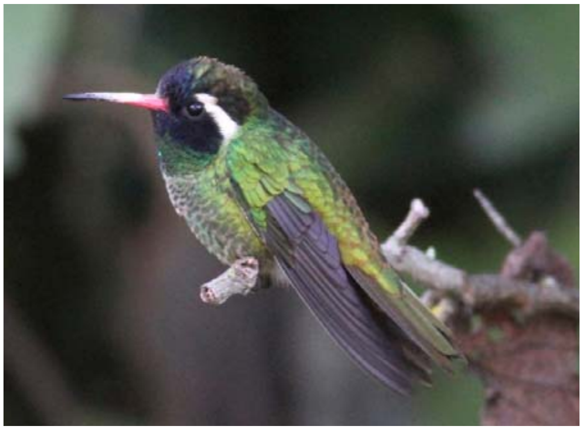

Definitive basic or formative male. Formative males reach full definitive appearance such as this following complete preformative molts and cannot be distinguished from definitive basic males. 13 Jan 2016, Jalisco, ML31576911.

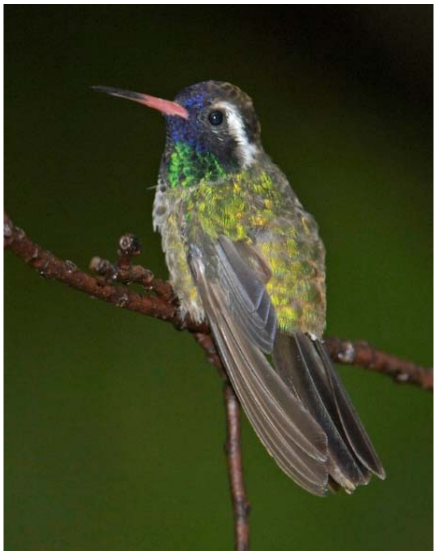

Male undergoing second or definitive prebasic molt. P6-p7 are dropped. 30 Jul 2006, Arizona, ML223535491.

**Figure S6. Images exemplifying molts and plumages in Violet-crowned Hummingbird.**

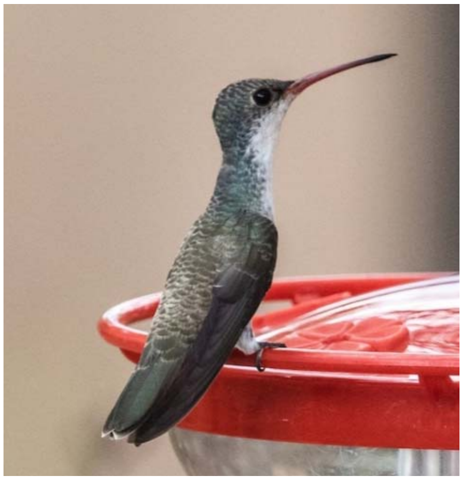

Juvenile. Note fresh scaly plumage, dull bill, and lack of purple iridescent feathers in crown. 5 Jul 2020, Arizona, ML247770321.

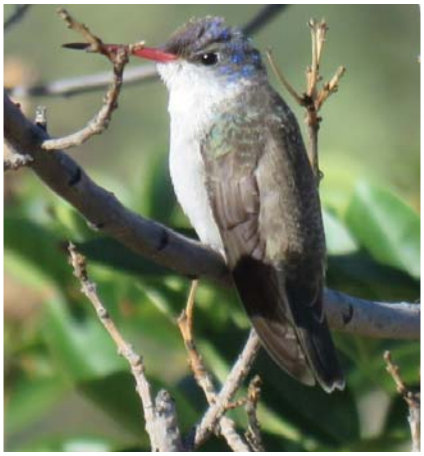

Individual undergoing preformative molt. Note fresh wings and brighter formative feathers in crown and among lesser coverts. Inner primaries can occasionally be replaced during this molt (Figure 3). 18 Nov 2017, Arizona, ML75461301.

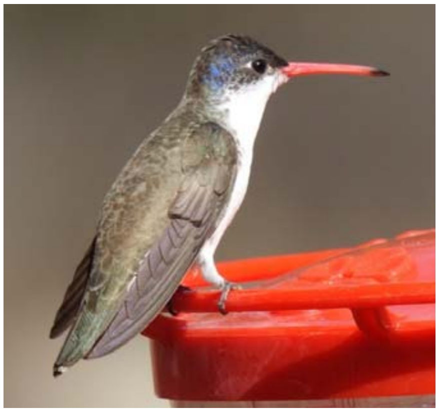

Formative plumage. Note retained brownish juvenile primaries and secondaries and molt limits in secondary coverts; crown is predefinitive in appearance. See also Figure 2. 27 Jan 2019, Arizona, ML137511331.

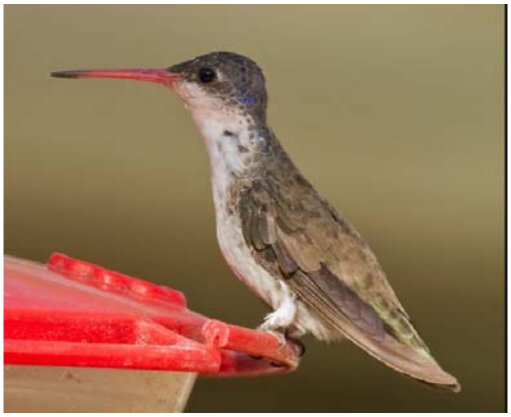

Individual completing second prebasic molt. Note unmolted juvenile p9-p10 and s3-s5; p8 is growing. Molt timing matches that of the second prebasic molt as the definitive prebasic molt only commences in May. 9 May 2011, Arizona, ML206043871.

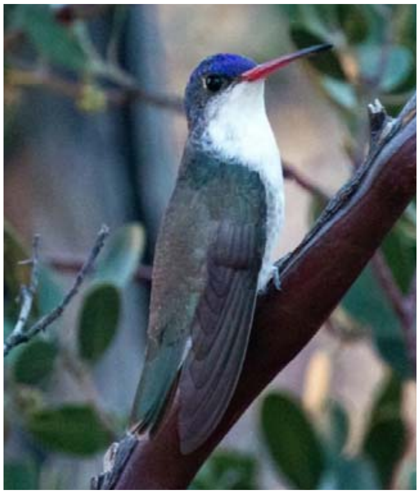

Definitive basic plumage. Note definitive appearance of body plumage, lack of molt limits in secondary coverts, and dusky remiges. 7 Jan 2020, Arizona, ML199909891.

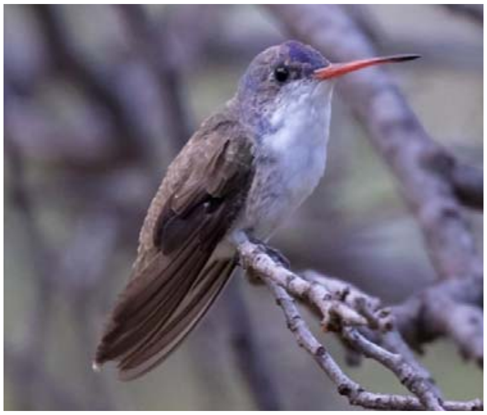

Male undergoing definitive prebasic molt. Note retained unmolted remiges are duskier than retained juvenile feathers. 10 Aug 2018, Arizona, ML111503391.

### Figure S7. Images exemplifying molts and plumages in Berylline Hummingbird.

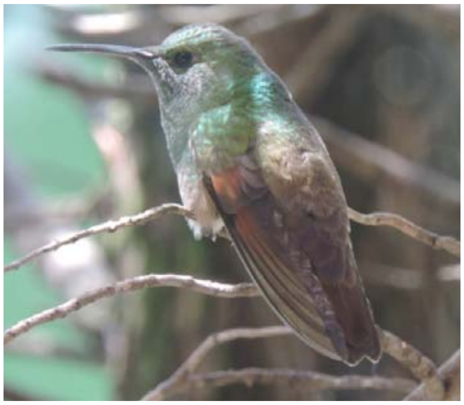

Juvenile. Note fresh scaly plumage, lacking glittering green. 27 Jun 2020, Distrito Federal, Mexico, ML246032741.

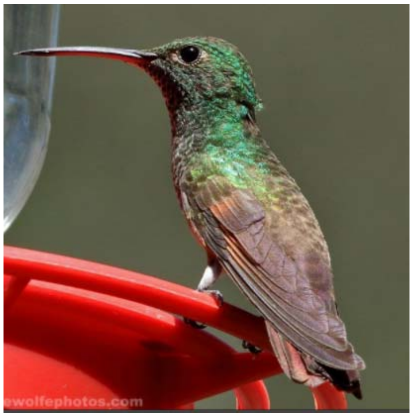

Individual undergoing preformative molt. Note mixed body feathering, brown remiges and greater coverts, and dull bill. 13 Aug 2019, Arizona, ML172361791.

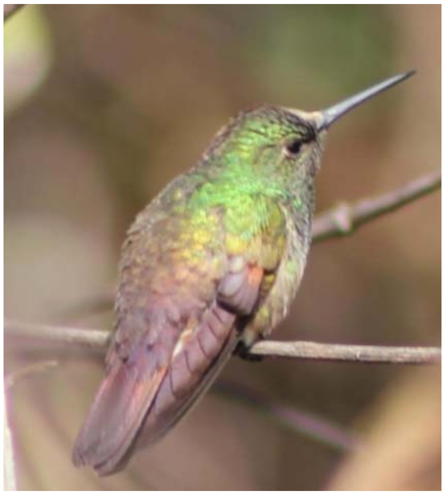

Formative plumage. Note mixed juvenile and formative body feathers and retained brown juvenile primaries and secondaries. See also Figure 2. 20 Jan 2018, Jalisco, Mexico, ML83111211.

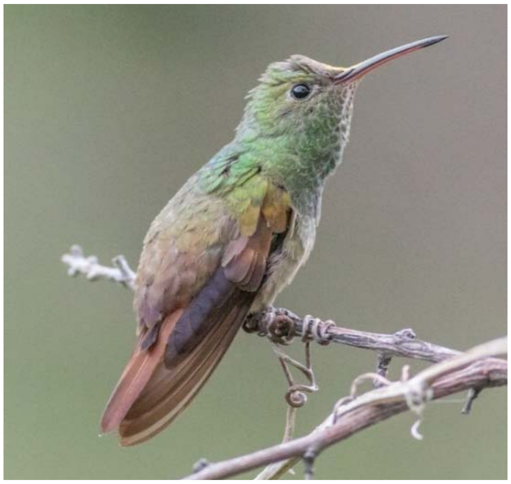

Individual undergoing second prebasic molt. Note unmolted juvenile p8-p10 and s2-s5; p7, s1, and s6 are growing. 5 May 2020, Arizona, ML233601741.

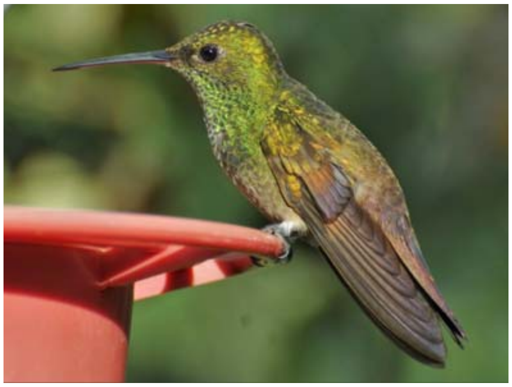

Definitive basic plumage. Note lustrous remiges and lack of molt limits among secondary coverts. 12 Aug 2009, Distrito Federal, Mexico, ML46645931.

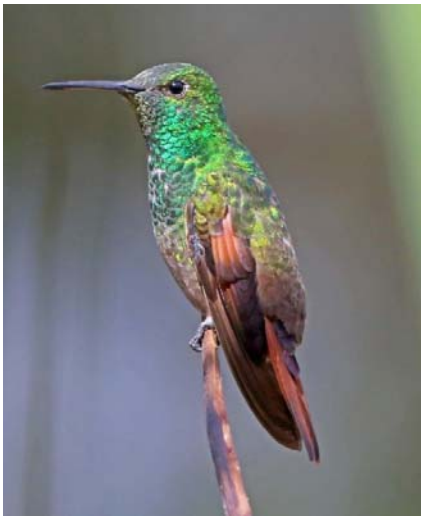

Individual undergoing definitive prebasic molt. Note unmolted outer primaries and secondaries are less worn than retained juvenile feathers. 10 Mar 2005, Distrito Federal, Mexico, ML46645931.

**Figure S8. Images exemplifying molts and plumages in Buff-bellied Hummingbird.**

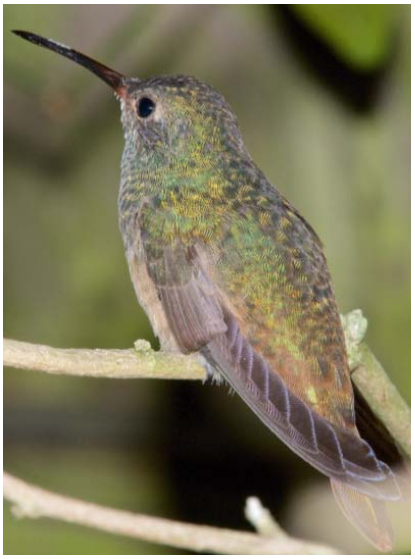

Juvenile. Note fresh scaly upperparts and lack of glittering feathers. 9 Mar 2017, Texas, ML50659971.

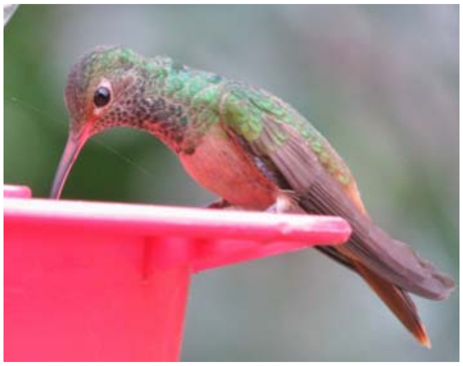

Individual undergoing preformative molt. Note fresh wings, brown juvenile remiges, and duller greater coverts. 8 Jan 2019, Texas, ML134258441.

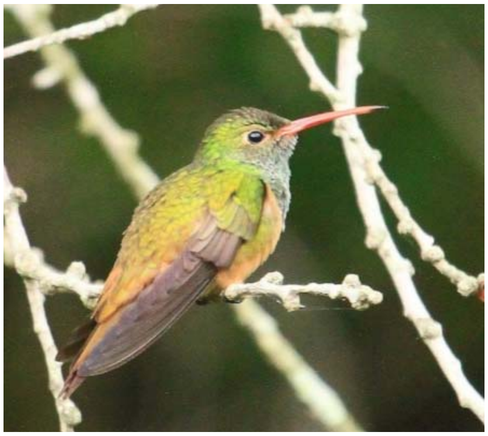

Formative plumage. Note retained brown juvenile primaries and secondaries and dull greater coverts. See also Figure 2. 18 Apr 2017, Texas, ML58643721.

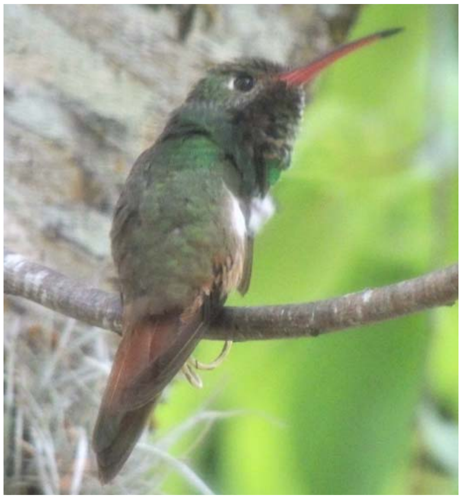

Individual undergoing second prebasic molt. Note unmolted juvenile p7-p10, outer rectrices, and most secondaries. 11 May 2017, Texas, ML57657391.

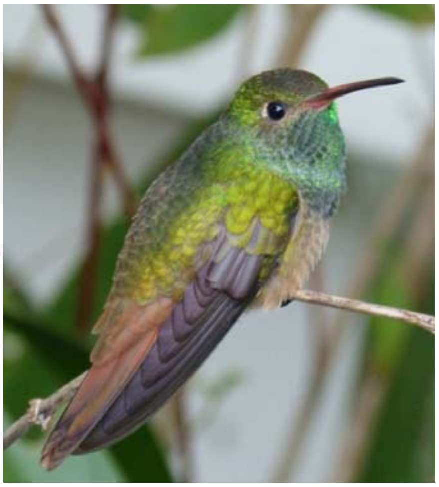

Definitive basic plumage. Note definitive body-feather appearance, lustrous dark remiges and basic rectrices. 26 Nov 2013, Louisiana, ML128861271.

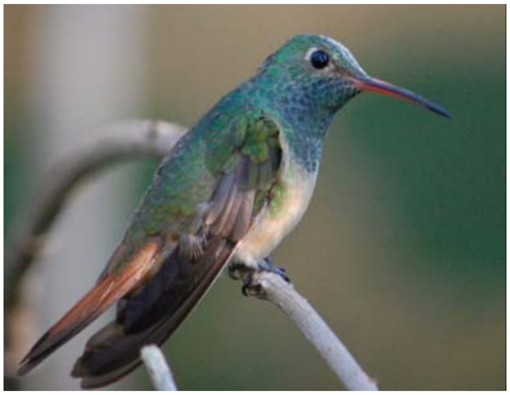

Individual undergoing definitive prebasic molt. Note definitive body-feather appearance, broad dusky unmolted outer primaries. 10 Nov 2018, Louisiana, ML122879681.

**Figures S9. Images of southwestern hummingbirds exemplifying ranges in variation of formative male plumages.**

Assessing the degree of definitive-like appearance in formative male hummingbirds can be challenging due to the effects of light and angle on iridescent gorget and other feathers. The following images were chosen as those that appear to depict the full extent of definitive-like appearance on each individual. When eBird checklist links are provided it indicates that several images of the same bird should be examined to fully appreciate extent of definitive-like appearance. Images represent worn formative birds during the period following the period of preformative molt but before commencement of the second prebasic molt. Five species are shown; not enough good images representing variation in formative Lucifer (*Calothrax lucifer*) and Berylline (*Saucerottia beryllina*) hummingbirds were available and for White-eared Hummingbird (*Basilinna leucotis*), formative plumage resembles definitive plumage in appearance and cannot be idenitfied in images.

Formative plumage is indicated in these five species by retained juvenile primaries, worn brown secondaries, and molt limits within the upperwing coverts. Links to Macaulay Library specimen pages (ML followed by 8 or 9 numerals) or eBird Checklists (S followed by 7 or 8 numerals) are provided and can be used to view each enlarged versions of each image, the photographer, and more detail on location and other aspects of the record. For four of these five species (all but Broad-billed Hummingbird), formative male plumage typically does achieve full definitive appearance.

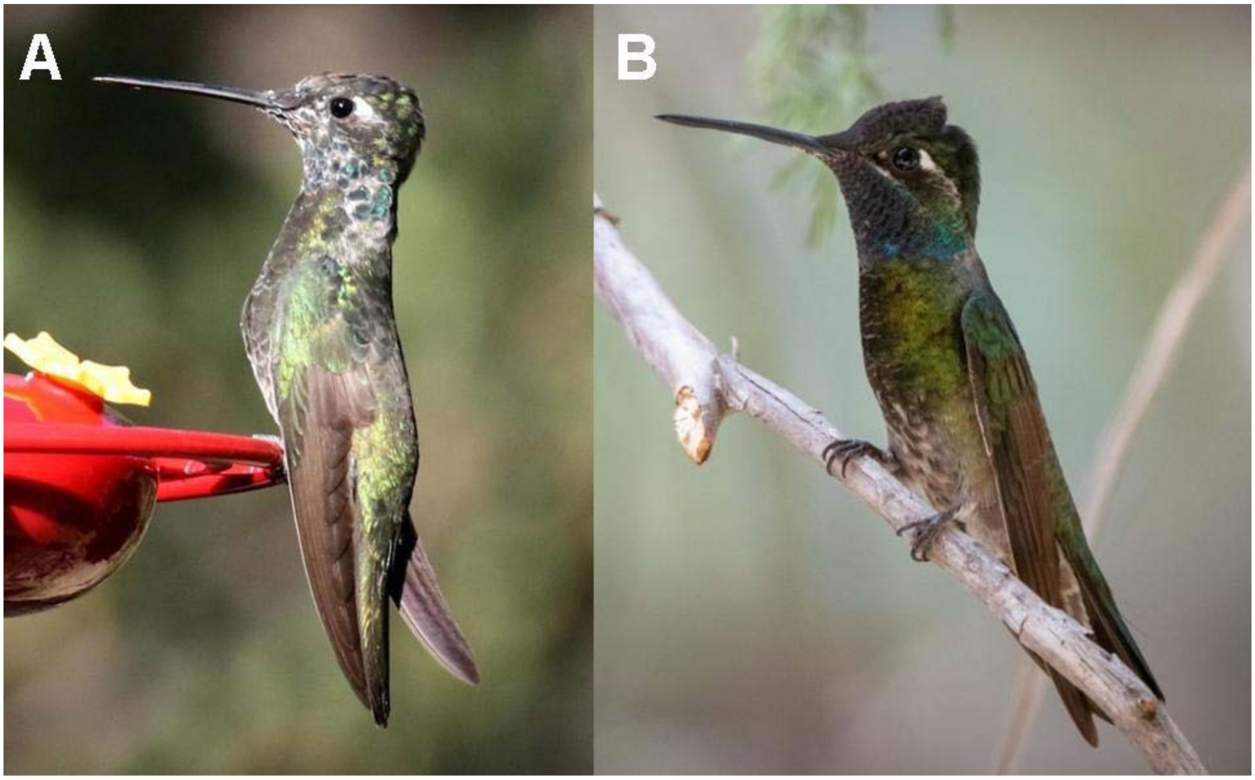

**Rivoli's Hummingbird (*Eugenes fulgens*)**. Minimal (**A**) and typical (**B**) development of definitive-like appearance in formative male plumage. Some definitive basic males can show brownish feathers in the belly as in B (though usually less); study needed whether or not this may be typical of second basic plumage. **A**: 23 May 2016, Arizona (ML29438681). **B**: 14 Jun 2020, Arizona (ML243435551).

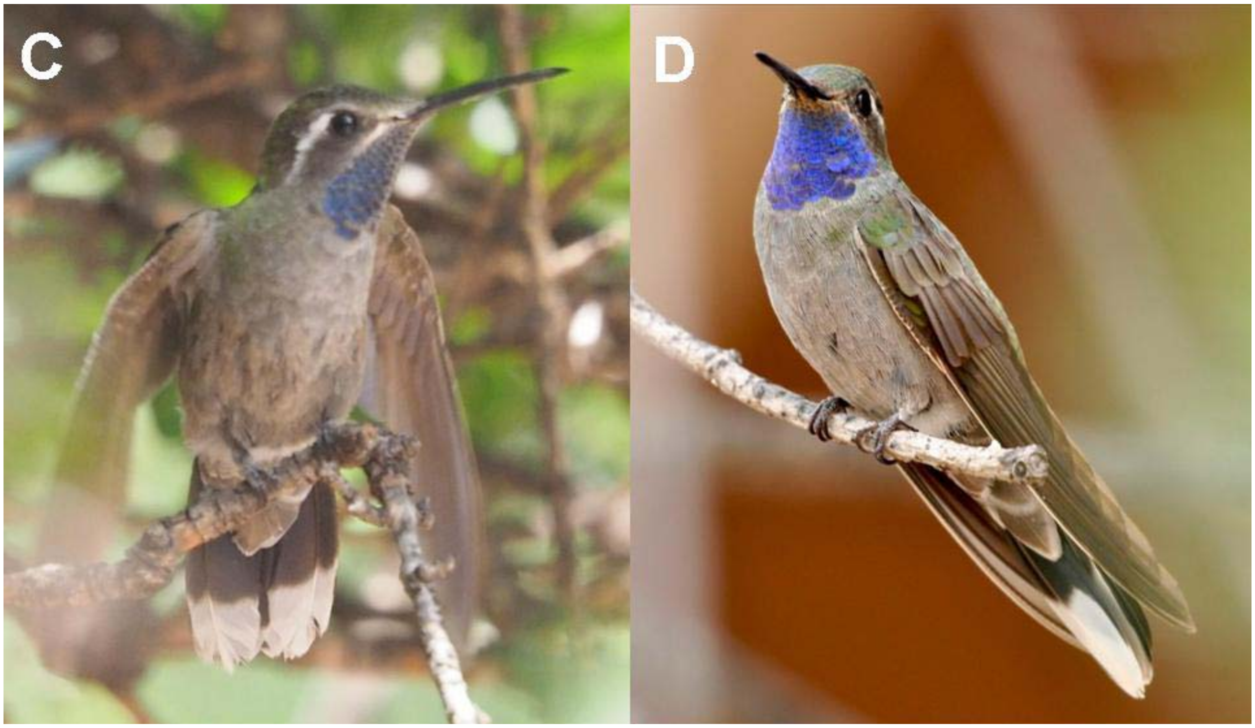

**Blue-throated Mountain-gem (*Lamphornis clemenciae*).** Minimal (**C**) and typical (**D**) development of definitive-like appearance in formative male plumage. None of 14 juveniles including nestlings (cf. S38554125) showed iridescent blue feathers, and no other hummingbirds show iridescent juvenile feathers, suggesting that these are formative but developed quickly during or following fledging (contra Pyle and Howell 2000). **C**: 10 Aug 2018, Arizona (S47766378). **D**: 14 Jul 2018, Arizona (S53611145).

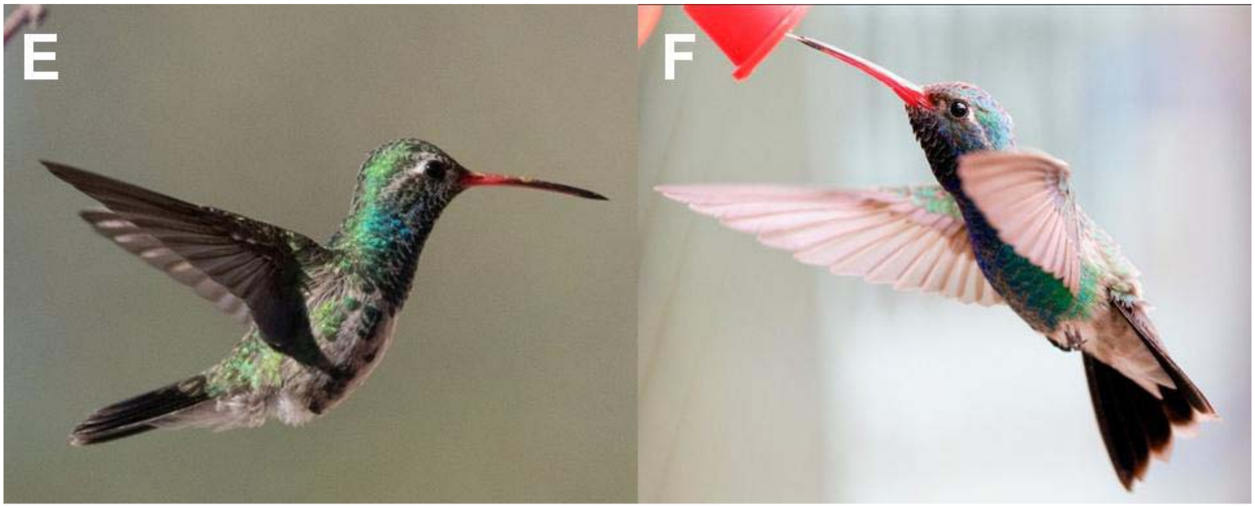

**Broad-billed Hummingbird (*Cynanthus latirostris*).** Minimal (**E**) and typical (**F**) development of definitive-like appearance in formative male plumage. The preformative molt and development of definitive-like appearance can vary from partial to complete (see also Figures 2, 3, and S4). **E**: 3 Apr 2020, Arizona (S66679154). **F**: 16 May 2019, Arizona (ML160635031).

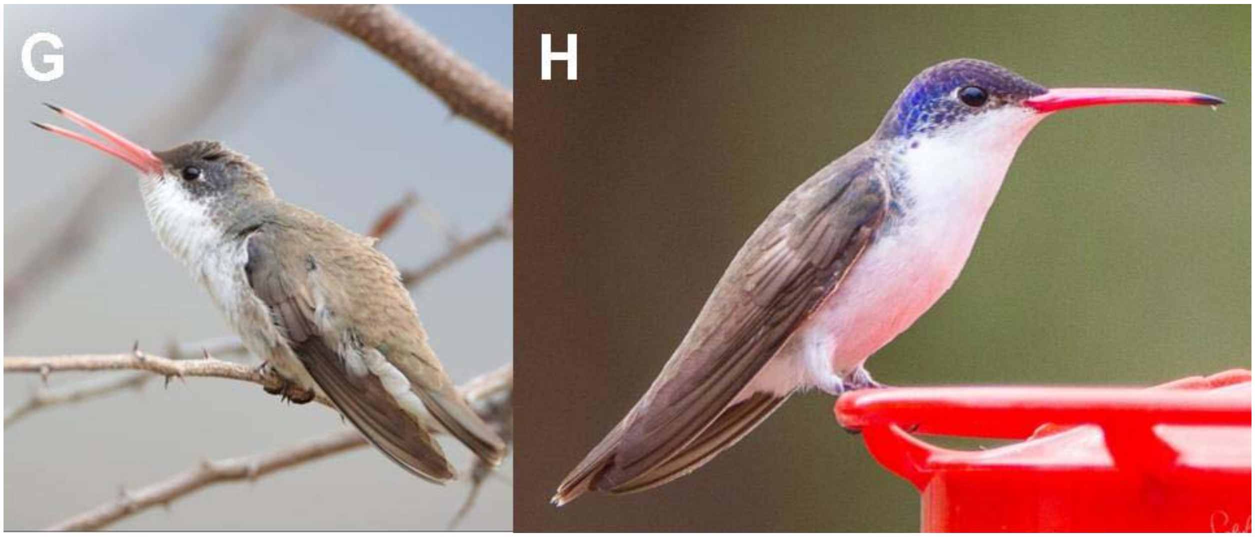

**Violet-crowned Hummingbird (*Leucolia violiceps*).** Typical (**G**) and maximum (**H**) development of definitive-like appearance in formative male plumage. See the eBird checklist for variation among photographs for the bird in **G**. **G**: 24 Mar 2018, Arizona (S43931142). **H**: 10 Mar 2018, Arizona (S43537351).

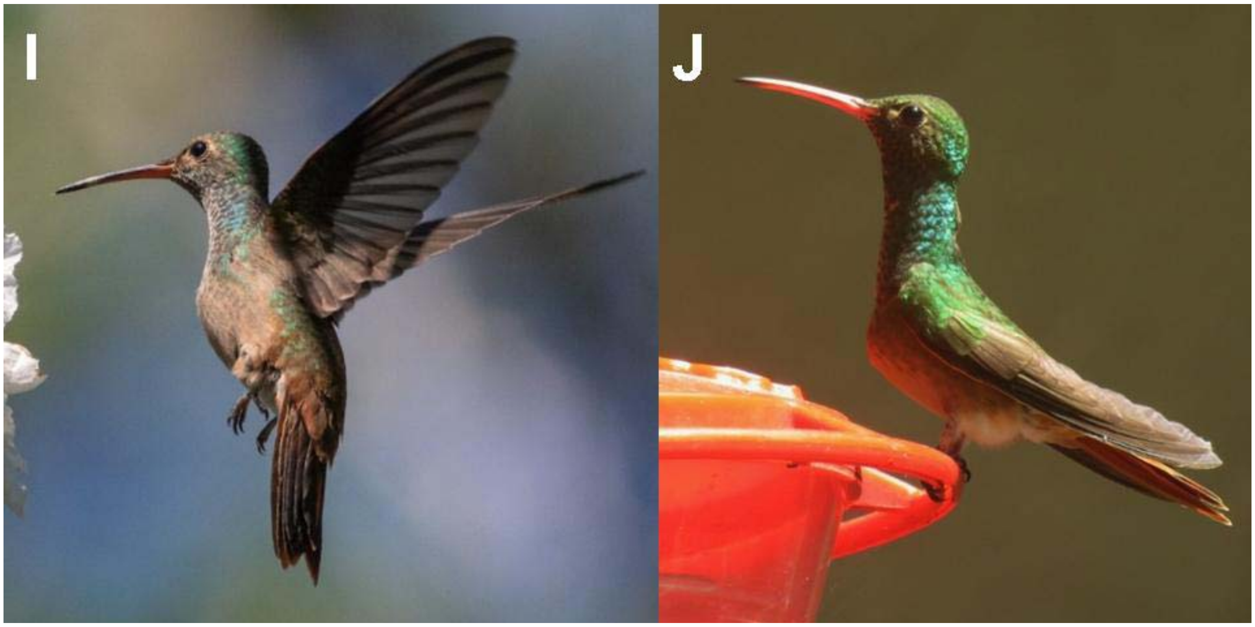

**Buff-bellied Hummingbird (*Amazilia yucatanensis*).** Minumum (**I**) and maximum (**J**) development of definitive-like appearance in formative male plumage. Formative plumage is found more widely throughout the year in Buff-bellied Hummingbirds than in the other southwestern species (Table S2). I: 19 Apr 2019, Texas (ML154689091). H: 28 Mar 2017, Texas (ML52660381).

**Appendix 1. Validation exercise undertaken by 17 participants including the author on 11 hummingbirds to assess whether or not field ornithologists can accurately collect data on avian molt from images.** (A) Participants were asked to rank their previous experience with banding and field observation as Low, Medium, or High, to fill out all cells for age, molt status, and condition of each primary (p1 to p10), and to estimate how many minutes it took to complete each line of data. "Correct" answers are given, those indicating the best possible conclusions by the author after evaluating all results. (B) Indicates the proportion of "correct" answers for each cell recorded by the 17 participants.

**Evaluation**

Participants correctly aged the 11 hummingbirds a mean 83% of the time, reached a correct conclusion on molt status 93% of the time, and provided correct answers for the condition of each primary from 83 to 91% of the time. For individual primary cells, correct answers ranged from 6% and 18%, up to 100% for most, with a mean of 87.2%. The mean time it took to complete each line was 3.7 minutes. The mean proportion of correct answers for the 132 cells was 87.5%. For the 17 observers this ranged from 80.3% to 95.4%. Among participants with Low, Medium, and High experience levels, correct answers were provided for 83.1% (*n*=3), 87.1% (*n*=3), and 88.9% (*n*=11) for banding experience and 87.6% (*n*=3), 86.6% (*n*=8), and 88.9% (*n*=6) for field experience, respectively.

There were no real patterns among errors on age determination, although most participants aged males in formative plumage and adult males in full definitive appearance correctly. For the three hummingbirds that were not in active molt (images 2, 6, and 10), correct answers were give by all 17 participants for molt status and for each primary condition. Some participants did not notice that the inner primaries had dropped for the birds in images 5, 7, and 11 and considered molt "inactive" for these. Several of the discrepancies for primary condition of individual feathers involved whether a primary had completed growth (score N) or not (score G). In image 9, for example, the author and others scored p6 as "N" but careful comparison with spacing of these primaries on other hummingbirds indicates that it is not quite fully grown, and should be scored "G," as some participants did. This is not a significant error, or one that would matter in molt analyses. For the White-eared Hummingbird in image 3, participants had trouble determining if the erupting pin feather was p5 or was p4, with p5 being missing. This resulted in low proportions of correct answers for the inner primaries, whether or not the pin feather was considered p4 or p5.

The biggest error rates occurred with the outer four primaries in images 1 and 4. Many participants did not take into account the sequence of outer primary replacement in hummingbirds, p8-p10-p9, and thus mistook the single remaining old primary as p10 rather than p9. In image 4, for example, p8 is "G," p9 is "O," and p10 is X. Only two participants recorded all three of these primaries correctly. In addition, many coded p7 on this bird as "N" but it is not yet completed growth and should be coded "G." In image 1, it that there are two old primaries left but one of these is the old primary from the other wing and difficulty in counting the inner primaries resulted in each of p7 and p8 being variously scored as N, G, or X. After Evaluation of inner primary spacing in other hummingbirds indicates that p2 is there but hard to see, and so the correct answer is P7 = N, p8 = G (not quite fully grown), p9 = O, and p10 = X. Only one participant scored the primaries on this bird correctly.

**A) Final Determinations for Hummingbird Exercise**

Age: Enter FY (first year, or HY/SY, through the 1st molt of primaries) or AD (adult, AHY/ASY, older)
Molt Status: Enter "active" if primaries are in active molt or "inactive" if not.
p1 to p10: Fill out only if you entered "active" for molt status. Enter N = New, X = not visible due to molt, G = visible and growing, O = Old
Minutes: Enter the number of minutes it took you to come to your conclusion, to the nearest 5 minutes.

| IMAGE | SPECIES | ML LINK | AGE | MOLT | P1 | P2 | P3 | P4 | P5 | P6 | P7 | P8 | P9 | P10 | MINUTES |
|---|---|---|---|---|---|---|---|---|---|---|---|---|---|---|---|
| 1 | RIHU | ML112128831 | FY | active | N | N | N | N | N | N | N | G | O | X | |
| 2 | VCHU | ML161385411 | AD | inactive | | | | | | | | | | | |
| 3 | WEHU | ML255139331 | FY | active | N | N | G | G | X | O | O | O | O | O | |
| 4 | BEHU | ML238319331 | FY | active | N | N | N | N | N | N | G | G | O | X | |
| 5 | BBIH | ML251781011 | AD | active | N | N | X | O | O | O | O | O | O | O | |
| 6 | BTMG | ML53554351 | FY | inactive | | | | | | | | | | | |
| 7 | BBIH | ML66989701 | FY | active | X | X | X | O | O | O | O | O | O | O | |
| 8 | RIHU | ML181183661 | AD | active | N | N | N | N | N | X | O | O | O | O | |
| 9 | BEHU | ML212588541 | FY | active | N | N | N | N | N | G | X | O | O | O | |
| 10 | BBEH | ML56665951 | FY | inactive | | | | | | | | | | | |
| 11 | VCHU | ML54332331 | FY | active | G | X | X | X | O | O | O | O | O | O | |

**B) Proportions of correct answers (n = 17 participants)**

| IMAGE | SPECIES | ML LINK | AGE | MOLT | P1 | P2 | P3 | P4 | P5 | P6 | P7 | P8 | P9 | P10 | Mean # of MINUTES |
|---|---|---|---|---|---|---|---|---|---|---|---|---|---|---|---|
| 1 | RIHU | ML112128831 | 0.82 | 1.00 | 0.94 | 0.94 | 0.94 | 1.00 | 1.00 | 1.00 | 0.82 | 0.18 | 0.47 | 0.06 | 5.1 |
| 2 | VCHU | ML161385411 | 1.00 | 1.00 | 1.00 | 1.00 | 1.00 | 1.00 | 1.00 | 1.00 | 1.00 | 1.00 | 1.00 | 1.00 | 2.8 |
| 3 | WEHU | ML255139331 | 1.00 | 1.00 | 0.71 | 0.82 | 0.35 | 0.59 | 0.59 | 1.00 | 1.00 | 1.00 | 1.00 | 1.00 | 3.2 |
| 4 | BEHU | ML238319331 | 0.71 | 1.00 | 1.00 | 1.00 | 1.00 | 1.00 | 0.88 | 0.88 | 0.18 | 0.35 | 0.35 | 0.35 | 4.1 |
| 5 | BBIH | ML251781011 | 1.00 | 0.77 | 1.00 | 0.82 | 0.71 | 0.71 | 1.00 | 1.00 | 1.00 | 1.00 | 1.00 | 1.00 | 3.2 |
| 6 | BTMG | ML53554351 | 0.82 | 1.00 | 1.00 | 1.00 | 1.00 | 1.00 | 1.00 | 1.00 | 1.00 | 1.00 | 1.00 | 1.00 | 3.5 |
| 7 | BBIH | ML66989701 | 0.71 | 0.71 | 0.65 | 0.65 | 0.59 | 1.00 | 1.00 | 1.00 | 1.00 | 1.00 | 1.00 | 1.00 | 3.9 |
| 8 | RIHU | ML181183661 | 0.71 | 0.94 | 1.00 | 1.00 | 1.00 | 0.88 | 0.71 | 0.71 | 0.82 | 1.00 | 1.00 | 1.00 | 3.5 |
| 9 | BEHU | ML212588541 | 0.82 | 0.94 | 1.00 | 1.00 | 1.00 | 1.00 | 1.00 | 0.12 | 0.65 | 0.65 | 1.00 | 1.00 | 4.0 |
| 10 | BBEH | ML56665951 | 0.77 | 1.00 | 1.00 | 1.00 | 1.00 | 1.00 | 1.00 | 1.00 | 1.00 | 1.00 | 1.00 | 1.00 | 3.1 |
| 11 | VCHU | ML54332331 | 0.82 | 0.82 | 0.12 | 0.77 | 0.77 | 0.59 | 0.65 | 1.00 | 1.00 | 1.00 | 1.00 | 1.00 | 3.9 |
| | **MEAN** | | **0.83** | **0.93** | **0.86** | **0.91** | **0.85** | **0.89** | **0.89** | **0.88** | **0.86** | **0.83** | **0.89** | **0.86** | **3.7** |

### ACKNOWLEDGMENTS

Foremost I thank the thousands of citizen scientists who have contributed images to the Macaulay Library and for agreeing to the license allowing use for research purposes. A total of 174 contributors provided images that are shown or linked in this study. The following photographers contributed to the Supplemental Materials section (see links to images for more detail). **For Supplemental Table S2:** images © Dale Adams, Jonathan Juárez Aguilar, Lisa Anderson, Juan Miguel Artigas Azas, Joe Baldwin, RJ Baltierra, Cathy Beck, Mikael Behrens, Tom Benson, Moe Bertrand, James Bozeman, Vicki Buchwald, Winston Caillouet, Nicola Cendron, Paul Conover, Greg Cook, Troy Corman, Ken Cox, Michael Crouse, Judith Ellyson, Richard Fray, Manuel Becerril González, Douglas Hall, Laurens Halsey, Hansel Herrera, Nancy Hetrick, Marla Hibbitts, Bill Hill, Sue Kaehler, Glenn Kincaid, Nick A. Komar Jr., Sergio Leyva, Michael Linz, Paul Maury, Christoph Moning, Peter Herstein, Rosie Howard, Tim Lenz, Max Leibowitz, Benjamin Miller, Shawn Miller, Randy Morgan, Brennan Mulrooney, Tina Nauman, Alan Van Norman, Carolyn Ohl, Bryant Olsen, Sig Olsen, Jen Owen, Richard Park, Paul Prappas, Donna Pomeroy, Vidal Prado, Kelly Rishor, Rose Ann Rowlett, Laval Roy, Shelley Rutkin, Ray Scally, Matthew Schmahl, Tim Schreckengost, Mel Senac, Jeff Sexton, Owen Sinkus, Larry Sirvio, Letha Slagle, Jim Stasz, Harlan Stewart, Jarrod Swackhamer, Rick Taylor, Daniel Garza Tobón, Jim Tonkinson, Rusty Trump, Eric VanderWerf, Ann Vaughan, Christopher Vincent, Nigel Voaden, Vincent Weber, Casey Weissburg, Sheri Williamson, Roger Woodruff. **For Figures S1-S8:** images © Ron Batie, James W. Beck, Moe Bertrand, Ernie Bradley, Paul Budde, Nicola Cendron, David Disher, Adam Dudley, Becca Engdahl, Allee Forsberg, Lisa Cancade Hackett, Thomas Ford-Hutchinson, Jim Gain, Manuel Becerril González, David Hollie, Brian Johnson, Tom Johnson, Eric Kallen, Frank Mantlik, Dan Maxwell, Suzie McCann, Michael McCloy, Mary McSparen, Dan Murphy, Marky Mutchler, Carolyn Ohl, Richard Park, Oliver Patrick, Lars Petersson, Edward Plumer, Carl Poldrack, Janet Rathjen, Phil Richardson, Arlene Ripley, Julio Alejandro Alvarez Ruiz, Brad Sale, Michael Schall, Bill Schmoker, Rosemary Seidler, Jeff Sexton, Larry Sirvio, Terry Sohl, Hans Spiecker, Susan Teefy, Ann Vaughan, Nigel Voaden, Ashley Wahlberg, Russ Wigh, and Steve Wolfe. **For Supplemental Figure s9:** images © Aaron Boone, Petra Clayton, Merryl Edelstein, Keith Gregoire, Mary McSparen, Oliver Patrick, Nick Pulcinella, Zach Skubiszewski, Lindsay Story, Mark Syvertson, Lyndie Mason Warner. See Acknowledgements in the primary manuscript for the names of additional contributors.
